# Interacting effects of cold snaps, rain, and agriculture on the fledging success of a declining aerial insectivore

**DOI:** 10.1101/2021.06.29.450344

**Authors:** Daniel R. Garrett, Fanie Pelletier, Dany Garant, Marc Bélisle

**Affiliations:** Département de biologie, Université de Sherbrooke, 2500 Boulevard de l’Université, Sherbrooke, Québec J1K2R1 Canada

**Keywords:** aerial insectivores, agricultural intensification, climate change, fledging success, precipitation, severe weather events, *Tachycineta bicolor*, Tree Swallows

## Abstract

Climate change predicts the increased frequency, duration, and intensity of inclement weather periods, such as unseasonably low temperatures (i.e., cold snaps) and prolonged precipitation. Many migratory species have advanced the phenology of important life history stages, and as a result will likely be exposed to these periods of inclement spring weather more often, thus risking reduced fitness and population growth. For declining avian species, including aerial insectivores, anthropogenic landscape changes such as agricultural intensification are another driver of population declines. These landscape changes may affect the foraging ability of food provisioning parents, and reduce the survival of nestlings exposed to inclement weather, through for example pesticide exposure impairing thermoregulation and punctual anorexia. Breeding in agro-intensive landscapes may thus exacerbate the negative effects of inclement weather under climate change. We observed that a significant reduction in the availability of insect prey occurred when daily maximum temperatures fell below 18.3°C, and thereby defined any day where the maximum temperature fell below this value as a day witnessing a cold snap. We then combined daily information on the occurrence of cold snaps and measures of precipitation to assess their impact on the fledging success of Tree Swallows *(Tachycineta bicolor)* occupying a nest box system placed across a gradient of agricultural intensification. Estimated fledging success of this declining aerial insectivore was 36.2% lower for broods experiencing four cold snap days during the 12 days post hatching period versus broods experiencing none, and this relationship was worsened when facing more precipitation. We further found that the overall negative effects of a brood experiencing periods of inclement weather was exacerbated in more agro-intensive landscapes. Our results indicate that two of the primary hypothesized drivers of many avian population declines may interact to further increase the rate of declines in certain landscape contexts.

## Introduction

Several avian groups are experiencing dramatic population declines (Rosenberg et al. 2019). Population estimates of many North American farmland and grassland birds, including aerial insectivores, suggest declines of over 50% since the 1970s (Stanton et al. 2018, Rosenberg et al. 2019). The spatio-temporal occurrence of these declines has led to the hypothesis that they are driven by a combination of global climate change and large-scale landscape modifications (Stanton et al. 2018, Spiller and Dettmers 2019), and particularly the process of agricultural intensification (Stanton et al. 2018).

In the northern hemisphere, global climate change has resulted in earlier spring temperatures (Hartmann et al. 2013). For instance, in USA the first fall frost has advanced by nearly 5.5 days and the last spring frost occurs nearly 7 days sooner when compared to the previous century (Kukal and Irmak 2018). The swift rate of change in timing of temperatures may lead to migrating species being unable to compensate for changes in thermally suitable habitats through dispersal, plasticity, or evolution (Crick 2004, Visser and Gienapp 2019). The inability of species to respond to such changes may generate or accentuate phenological mismatches between peaks of seasonal food resources and peaks in resource requirements of migratory species, and as a result, cause fitness decreases due to overall mistimed breeding schedules (Visser and Gienapp 2019). Furthermore, global climate change predicts increases in the frequency and intensity of inclement spring weather, such as swift changes in ambient temperature (e.g., cold snaps) and prolonged periods of precipitation (Rahmstorf and Coumou 2011, Wuebbles et al. 2014). Though often difficult to define (van de Pol et al. 2017), extremes in weather (e.g., storms, blizzards, and oceanic heatwaves) can have significant impacts on ecosystem functions (reviewed in Ummenhofer and Meehl 2017). In birds, even slightly lower than average daily temperatures or greater than average precipitation can lower fitness (Tuero et al. 2018, de Zwaan et al. 2019, Skwarska et al. 2021). Individuals experiencing such periods of inclement weather may be subjected to the reduction of temperature-dependent food resources, as well as to the additional consequence of the direct effects of poor weather (Pipoly et al. 2013, Moreno et al. 2015, Arbeiter et al. 2016, Marcelino et al. 2020). Therefore, migrating species adapting to global changes through advancing their spring migration, may be at greater risk of experiencing periods of inclement weather (Both et al. 2010, Visser and Gienapp 2019). The phenological advancement of several declining species has recently been observed (Dunn and Winkler 1999, Møller et al. 2006, Townsend et al. 2013, Bourret et al. 2015). For example, two Tree Swallow (*Tachycineta bicolor*) populations located respectively in Ontario, Canada, and New York State, USA, exhibit both chronic population declines and advanced clutch initiation dates (Dunn and Winkler 1999, Shutler et al. 2012). During the same period of declines, a significant increase in the number of poor weather days (e.g., unseasonably low temperatures or prolonged precipitation) was observed and resulted in elevated nestling mortality (Cox et al. 2019, Shipley et al. 2020).

The declines of farmland and grassland birds are correlated with not only climate change but also with the process of agricultural intensification (Stanton et al. 2018, Spiller and Dettmers 2019). Hypothesized agricultural drivers of declines are divided into direct (e.g., habitat loss, mechanization resulting in nest destruction, or acute pesticide exposure) and indirect effects, principally through the reduction in prey availability or sublethal pesticide exposure (Benton et al. 2003, Stanton et al. 2018). Certain agricultural practices are observed to alter insect populations and communities through changes in abundance, phenology, or species composition and interactions (Grüebler et al. 2008, Pisa et al. 2015, Wagner 2020). Insectivorous birds breeding in modern agricultural landscapes, including aerial insectivores, could thereby be subjected to reduced availability of prey items (Poulin et al. 2010, Nocera et al. 2012, Garrett et al. 2021a).

Agricultural drivers of population declines may also impact the physiology of birds in ways that influence their capacity to respond adaptively to periods of inclement weather. For instance, sublethal exposure to organophosphate and carbamate insecticides may induce short-lived hypothermia, likely due to the impairment of thermoregulation (Grue et al. 1997). Moreover, sublethal exposure to neonicotinoid insecticides may result in anorexia (Eng et al. 2019), thereby aggravating reduced thermoregulatory capacity and other challenges posed by reduced prey availability (Garrett et al. 2021a). Birds breeding within more agro-intensive landscapes are at elevated risk of exposing themselves or their offspring to such agents (DiBartolomeis et al. 2019, Malaj et al. 2020, Sigouin et al. 2021) and may subsequently be less likely to survive through periods of poor weather. Furthermore, landscapes dominated by row crop monocultures express landscape simplification in which large swaths of areas are occupied by only a handful of habitats (Benton et al. 2003). This phenomenon may make finding and exploiting suitable food resources more difficult for animals relying upon “residual” marginal habitats, as landscape simplification lowers the functional connectivity of agricultural landscapes (Hinsley 2000, Bélisle 2005, Rainho and Palmeirim 2011). Parents may compensate for poor breeding landscape quality by increasing foraging effort, but such increases may result in reduced body condition (Hinsley 2000, Olsson et al. 2008). Periods of inclement weather may thus exacerbate the added stressors stemming from breeding within more agro-intensive landscapes (Stanton et al. 2016, Staggenborg et al. 2017, Evens et al. 2018, Garrett et al. 2021b), and may even influence the capacity for parental care and investment (de Zwaan et al. 2019). Therefore, given that selection favors earlier breeding events (Sheldon et al. 2003, Porlier et al. 2012, Marrot et al. 2017), it seems imperative to evaluate how the expected increase of inclement weather events may interact with the consequences of breeding within more agro-intensive landscapes.

Here we present the results of an 11-year study monitoring the breeding success of a Tree Swallow population experiencing a wide range of spring temperatures and precipitation. This population breeds within a nesting box system placed along a gradient of agricultural intensification in southern Québec, Canada. We first identified a critical temperature in which the availability of their main prey, namely Diptera (Bellavance et al. 2018), changed significantly, using an approach similar to Winkler et al. (2013). We then defined days in which ambient temperature fell below this critical temperature as a day representing a cold snap. We then evaluated the interacting roles that cold snaps, prolonged precipitation and the gradient of agricultural intensification had on fledging success within this population. As observed by Cox et al. (2019) and Shipley et al. (2020), we expected cold snaps to reduce fledging success, and the effect size to increase with both the increasing number of cold snaps and the total amount of precipitation experienced by broods. Finally, we expected the severity of these relationships to increase as breeding landscapes became increasingly composed of agro-intensive cultures.

## Methods

### Study area and nest box system

In 2004, 10 nest boxes, spaced approximately 50 m apart, were placed along the field margins of 40 separate farms dispersed across a gradient of agricultural intensification in southern Québec, Canada (Fig. 1; see Ghilain and Bélisle 2008 for details). In 2006 we started intensively monitoring the breeding activity of these nest boxes and continued through the breeding season of 2016. This monitoring protocol resulted in 400 nest boxes being actively monitored throughout this period. The gradient of agricultural intensification was characterized by an eastwest shift of agricultural production. The eastern portion of the system was composed primarily of pastures and forage crops (e.g., hay, alfalfa *(Madicago sativa),* and clover *(Trifolium spp.*)), requiring less agricultural inputs and interspersed with large expanses of forest cover. The west was dominated by large-scale row-crop monocultures (principally corn (*Zea mays*), soybean *(Glycine max)* and wheat *(Triticum spp.))* and was denuded of forest cover (Jobin et al. 2003, Ruiz and Domon 2009). Increased use of monocultures has resulted in a near reliance on fertilizers, pesticides, mechanization and drainage of surface waters and wetlands (Jobin et al. 2003). Between 2011 to 2019, nearly 100% of the corn and 60% of the soybean were sown as neonicotinoid-coated seeds in our study area (MDDELCC 2015). As a consequence, the rate at which several pesticides were detected above the levels deemed safe for the chronic exposure for aquatic life has increased in the surface waters of this region (Giroux 2019).

**Fig. 1:**
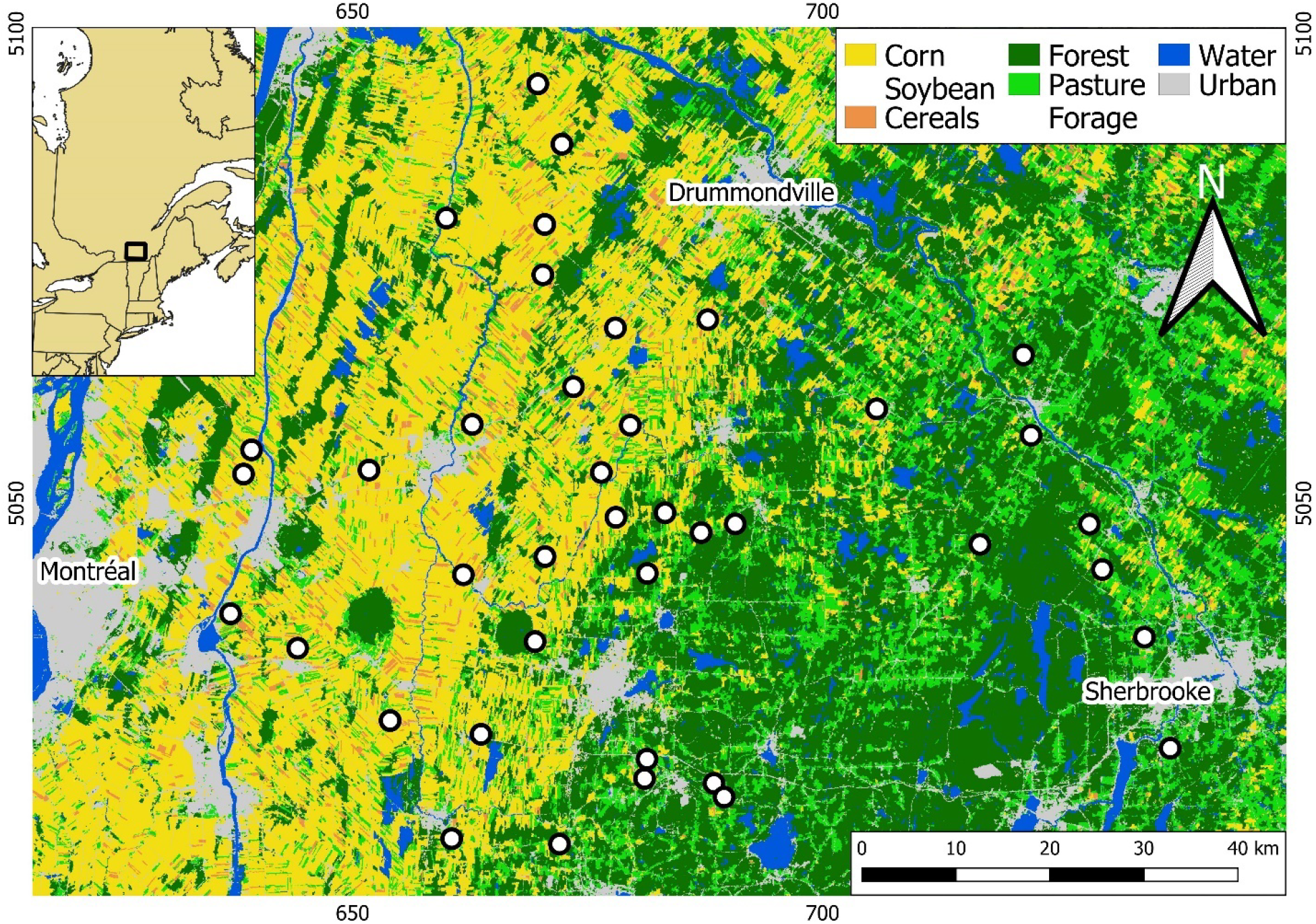
Location of the 40 farms (white dots) monitored across a gradient of agricultural intensification within southern Québec, Canada. Underlying image represents the agricultural gradient derived from the Annual crop inventory of 2013 (AAFC, 2018). Each pixel is either one of the five higher order land cover categories or open water. The box within inset map denotes the extent of the main map.

### Nest box monitoring

Starting prior to the arrival of Tree Swallows, we monitored the breeding activity in nest boxes every two days throughout each breeding season. This monitoring schedule allowed us to estimate the dates of laying, incubation, hatching, and fledging, as well as to record the number of eggs, nestlings, and fledglings of each breeding event. On average, first-clutch-occupancy by Tree Swallows was 60% ranging between 51% and 74% throughout the 11-year data set. We caught and banded adult females during incubation and adult males during food provisioning. We were 99% and 80% successful, respectively, at capturing targeted adults. We applied an aluminum US Geological Survey (USGS) band containing a unique identification code to unbanded adults upon capture, as individuals can breed within this system over multiple breeding seasons.

### Insect sampling and local prey availability

Throughout the study, 2 insect traps were placed on each of the 40 farms (N=80). Traps were spaced ~ 250 m apart along the central portion of each nest box transect. Trap content was collected every two days throughout each breeding season (see Garrett et al. 2021a for details). The processing of insect samples focused on the period between 1 June and 15 July, covering the nestling rearing period of 96% of the monitored breeding attempts. The content of traps included a wide range of insects stemming from many of the surrounding habitat types, and that overlap significantly the diet of nestlings (Laplante 2013, Bellavance et al. 2018). In this system, 74% of the biomass of the food boluses provided to nestlings were Diptera (Bellavance et al. 2018), a pattern shared with other Tree Swallows study systems (Winkler et al. 2020). We thus used the biomass of Diptera within each sample as a proxy of local prey availability. It should be noted that, in a separate analysis of fledging success using this same data set, models incorporating prey availability measures represented by Diptera outperformed ones in which prey availability was represented as all invertebrates (Appendix from Garrett et al. 2021a). Once extracted, Diptera samples were placed in an oven at 60°C for over 24 hours, ensuring no further change in biomass occurred and were then weighed without delay to the nearest 0.0001 g. The dried biomass of Diptera from these samples (hereafter referred to as Diptera biomass) thus represents an estimate of local prey availability on a specific farm and year over the two days prior to the sample collection date.

### Landscape composition

Landscape composition (i.e., the relative coverage of habitats composing a given landscape) focused on habitats influential to Tree Swallows and ones composing landscapes throughout our study system [e.g., tree cover, open habitats, and cover of surface water, (Rendell and Robertson 1989, 1990, Winkler et al. 2020)]. The relative cover of habitats was calculated within 500 m of nest boxes, a spatial scale in which ~80% of food provisioning adult female Tree Swallows spend their time (Elgin et al. 2020, Garrett et al. 2021b). Tree Swallows are often associated with open bodies of water and will travel distances greater than 500 m to forage over them (Elgin et al. 2020, Garrett et al. 2021b). The relative cover of this habitat within 500 m of nest boxes and in the agricultural contexts covered by this study is low (0.66% ± 1.07%). Recognizing this habitat as important to Tree Swallows (Berzins et al. 2021), we calculated its relative cover at 3 km from nest boxes following methods presented in Garrett et al. (2021a). Calculations used vector layers acquired from the Canadian National Hydro Network (NHN, 2020) and the sf (Pebesma 2018) and rgeos (Bivand and Rundel 2019) packages in R version 3.6.2 (R Core Team 2019). Characterization within 500 m of nest boxes occurred separately for each farm at the end of each breeding season to facilitate crop identification. Parcels representing different habitats and agricultural fields were first delineated using orthophotos (scale 1:40,000) in QGIS (version 3.16) (QGIS 2020), and then characterized *in situ.* We determined which cultures, if any, were in agricultural fields and then reclassified them into forested, corn and soybean, forage fields (including hay, other grasses, alfalfa, clover, and pastures), and cereals (principally wheat, and to a lesser extent, oat *(Avena spp.)* and barley *(Hordeum spp.)).* We then calculated the mean percent cover of these habitats across the 10 nest boxes on each farm and for each year independently.

We desired an integrative measure of the percent cover of all habitat groups observed for each year and farm combination, defined as the landscape context. We therefore used a robust principal components analysis (PCA) for compositional data on the entire landcover data set to assign “site scores” to each of the farms during each year (Filzmoser et al. 2009). Site scores were the year and farm specific values along the first two components of the resulting compositional PCA. The single PCA was fitted using the robCompositions package (Templ et al. 2011) in R, resulting in the calculation of 440 different landscape contexts (i.e., 40 farms x 11 years). Site scores were assigned to each breeding attempt and insect sample and used in all subsequent analyses.

The PCA’s first component (Comp.1) explained 80.34% of the variance in landcover, correlating positively with corn and soybean and negatively with both forage fields and forest cover. The second component (Comp.2) explained 14.69% and correlated negatively with forage fields and positively with forest cover (Fig. 2). To avoid overly complex models, we included only these two components to represent the landscape context. Landscapes expressed by minimizing Comp.1 and Comp.2 values represent ones for which there is an abundance of forage fields and above average forest cover and are referred to as forage landscapes.

**Fig. 2:**
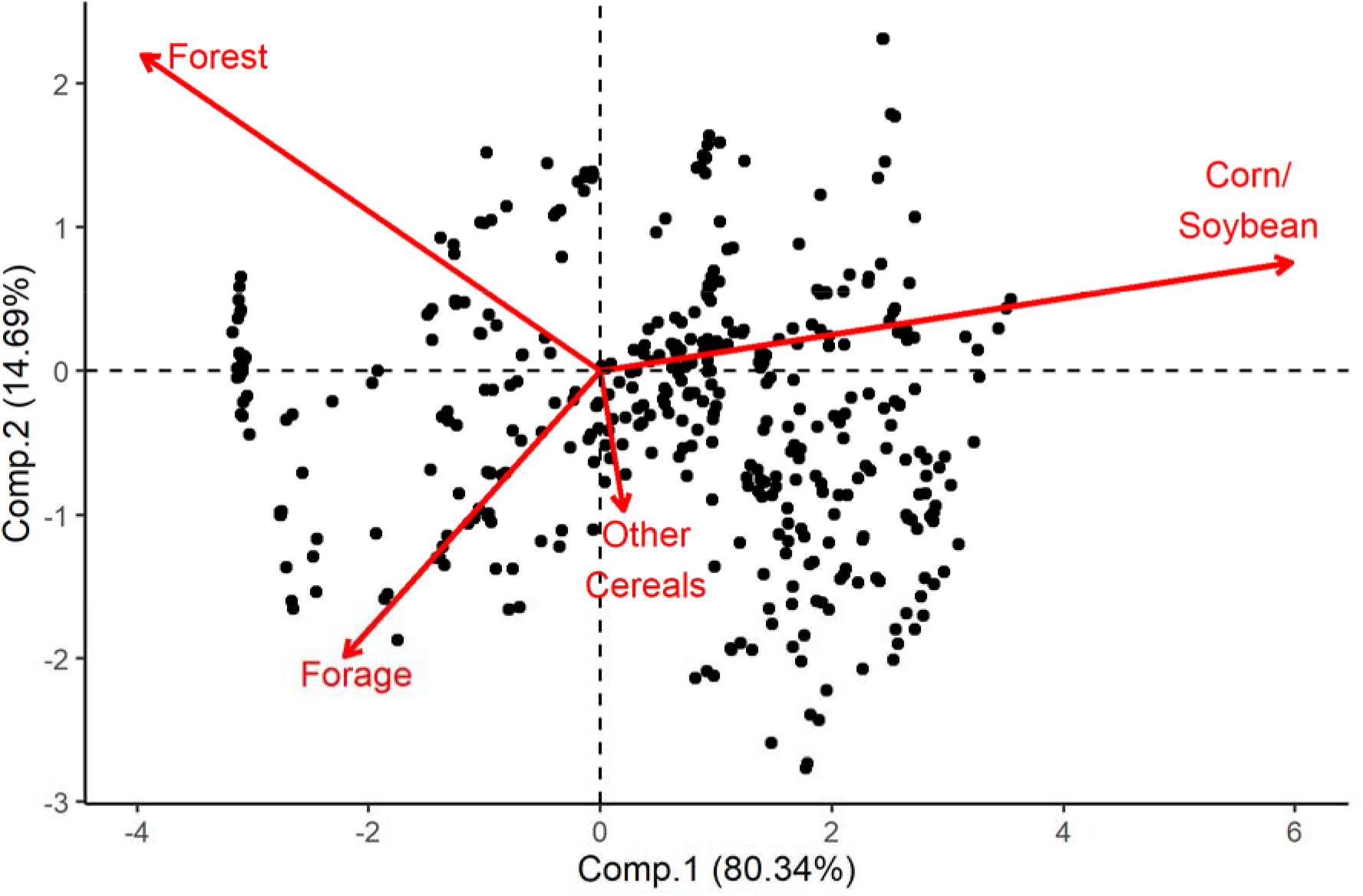
Robust compositional principal components analysis (PCA) of the landscape habitat composition surrounding each of the 40 farms monitored between 2006 and 2016. Arrows indicate the eigenvalue and loadings of each higher order land cover. Points within the background represent the site scores assigned to each farm and year combination and used to define the landscape context within 500 m of nest boxes (N=440 farms*years).

### Weather variables

We monitored hourly temperature with iButtons (model DS1922L; Embedded Data Systems, Lawrenceburg, Kentucky, USA) attached to the underside of a single nest box on each farm. Temperatures were recorded throughout each breeding season and started prior to the spring arrival of Tree Swallows. We derived daily summaries of the minimum, mean, and maximum daily temperatures between daytime hours (05:00 to 20:00). We measured total farm-specific precipitation every two days throughout the breeding season using a single pluviometer placed on each farm and recorded measurements to the nearest 0.5 ml during each farm visit.

### Statistical analyses

#### Cold-snap days

Cold snap values were grounded in the functional response of local prey availability to maximum daily temperature, as we expected insects to require a threshold temperature before they are active (Williams 1961, Grüebler et al. 2008, Roitberg and Mangel 2016). We calculated the mean maximum daily temperature between the day of and the day prior to sample collection (mean maximum daily temperature). We then used a similar approach to that of Winkler et al. (2013) using generalized additive mixed effects models (GAMMs) to model the relationship between Diptera biomass and mean maximum daily temperature to determine the temperature at which the instantaneous rate of biomass change was greatest. The cold snap threshold temperature was thus calculated as the temperature corresponding to the maximum of the first derivative of the relationship between Diptera biomass and mean maximum daily temperature. Peak rate in change was chosen because temperatures around this critical threshold likely result in dramatic differences in prey availability (Williams 1961, Roitberg and Mangel 2016). We undertook an extensive modeling exercise to determine the treatment of the functional relationship between these variables, including evaluating the hypothesis that the critical threshold temperature varies along the agricultural intensification gradient (full details in Appendix S1). Modeling used the “bam” function from the mgcv package in R (Wood 2015). Diptera biomass was modeled with a Gamma distribution with a log link function. The influence of mean maximum daily temperature was treated as a tensor product smooth, and in order to evaluate overall trends, we kept the basis dimension of this predictor low (k=10). In all models, we included the effect of Julian date of sample collection as a tensor product smooth and precipitation as a linear predictor to account for phenological variations in biomass. We further included the year and farm as random effects to control for the hierarchical structure of the sampling design (Pedersen et al. 2019). All correlations and pairwise functional responses are provided in Appendix S1 (Fig. S1).

#### Fledging success

We limited analyses of fledging success to first breeding attempts (90% of all attempts and ranging between 80% and 95% across years), as reproductive success varies greatly between first and second attempts, and second attempts occur near exclusively following the failure of a nest during laying or incubation (Robertson and Rendell 2001). The date at which insect samples started to be processed placed a constraint on the breeding data we could use. Most notably, very early breeding attempts were omitted due to hatching occurring prior to the start of the processing window (~1% omitted). It should be highlighted that during the 11-year data set presented, the likelihood of a brood being omitted because its hatching date occurred prior to insect sampling was irrespective of the breeding season. Fledging success was modeled as the ratio between the number of fledglings and the number of nestlings (i.e., number of successes over trials) via generalized linear mixed effects models (GLMMs) with a binomial distribution and logit link function. The year, farm, and nest box IDs were included as random factors to account for the hierarchical structure of the data set (i.e., nest box nested within farm nested within year). Model covariate summaries can be found in Table 1 and were averaged across a 12-day window post-hatching. This period is when Tree Swallow nestlings become homeothermic, reach peak body mass, and experience the greatest rate of mortality, thus representing the period where resource availability is presumably most crucial (McCarty and Winkler 1999, Houle et al. 2020). Cold temperatures and precipitation may act both directly on nestlings through thermoregulation and indirectly through a reduction in resource availability. To control for trophic mediated indirect effects of weather on fledging success, we derived estimates of local prey availability during the nestling rearing period of each breeding attempt. Prey availability was represented by predictions from generalized additive models (GAMs) in which raw values of Diptera biomass were regressed against the Julian date of sample collection for each farm and year separately. Modeled Diptera availability captures its general phenology throughout each season and avoids biases caused by more punctual or local disturbances (e.g., capturing an insect swarm). From these predictions, we calculated an estimate of the mean prey availability during the respective 12-day window of each breeding attempt. GAMs were fitted as a tensor product smoother and an identical degree of smoothness (k=10).

**Table 1:** Mean (±SD) of covariates used to model fledging success of 1,897 Tree Swallow broods between 2006 and 2016. Covariates were averaged over a 12-day period after hatching. Variable groups identify which covariates were present within models found in Appendix S2: Table S1.

| Covariate | Acronym | Unit | Mean ± SD | Variable group |
| --- | --- | --- | --- | --- |
| Site score Comp.1 | Comp.1 | - | 0.20 ± (1.80) | Base/Land |
| Site score Comp.2 | Comp.2 | - | -0.18 ± (0.91) | Base/Land |
| Cold snap days | CSD | (day) | 0.93 ± (1.14) | Base/Snap |
| Precipitation | RN | (ml) | 7.44 ± (4.92) | Base/Rain |
| Open water | WT | (%) | 0.01 ± (0.01) | Base |
| Prey availability | PA | (g) | 0.03 ± (0.02) | Base |
| Hatching day | HD | (Julian day) | 159.91 ± (5.76) | Base |
| Brood size | NN | (nestlings) | 4.83 ± (1.27) | Base |

We took an information theoretic and multimodel approach to assess how weather influenced fledging success and if its effect varied with landscape context (Burnham and Anderson 2002). We compared a set of competing models, including a Null model with only random effects. These data were previously used to determine the combined influence of prey availability and landscape context within our study area on fledging success, and as a result a most predictive and parsimonious model had then been determined (Garrett et al. 2021a). We thus compared all subsequent models to this base model (Base). This model included the brood size, age of the breeding female [second year (SY) vs. after second year (ASY); (Robertson and Rendell 2001)], hatching date, percent cover of open water within 3 km of the breeding attempt, the interaction between site scores (Comp.1 and Comp.2) and the prey availability estimate. To these variables we also added the number of cold snaps and the mean daily precipitation during the 12 days following hatching, including the interaction between these variables, as these factors are identified as influential to Tree Swallows from other study systems (Cox et al. 2019, Shipley et al. 2020). It should be explicitly stated that our measure of cold snap and precipitation occurrence cannot measure the consecutiveness or coincidence of these events. Instead, these measures represent a coarse proxy of the weather that broods experienced during this time window. All model terms within Base were included in all subsequent models of the candidate set. Principal interests were in the interactive effects of cold snaps, precipitation, and the gradient of agricultural intensification on fledging success. We included models with interaction terms between Comp.1 and the number of cold snaps (Base + Land*Snap), between Comp.1 and the mean precipitation (Base + Land*Rain), or both of these interaction terms (Base + Land*Snap + Land*Rain). We further predicted these factors may interact with one another and included a model with a three-way interaction between these three variables (Base + Land*Snap*Rain). A summary table of all continuous covariates can be seen in Table 1 and the correlations and functional relationships between all these variables are provided in Appendix S2 (Fig. S1).

It should further be noted that we explored our choice of temperature threshold and post hatching window in an extensive modeling exercise (Appendix S3). This modeling endeavor did not highlight any qualitative differences from the following results. We therefore continued to utilize the methodology presented thus far in an attempt to build models based on sound biological variables influential to the behavior and physiology of Tree Swallows (Burnham and Anderson 2002, Harrison et al. 2017).

In models of fledging success, the effects of key individual variables, including interactions, were estimated via multi-model inference whereby predictions were calculated by model-averaging with shrinkage and shown with their 95% unconditional confidence intervals (Burnham and Anderson 2002). All quantitative covariates were Z-transformed, ^-squares calculated following Nakagawa and Schielzeth (2013), and analyses performed in R using the glmmTMB (Brooks et al. 2017) and AICcmodavg (Mazerolle 2020) packages. Model validation, including evaluation of normally distributed residuals (simulated), heteroscedasticity, and collinearity checks with variance inflation factors, followed Zuur et al. (2009) and used the DHARMa package (Hartig 2020).

## Results

### Effects of temperature on local prey availability

We collected and processed 15,916 insect samples from 8,614 farm visits. Mean Diptera biomass (± SD) was 0.030 ± 0.044 g (per trap), ranging between 0.019 g and 0.037 g across years. Overall mean maximum daily temperature (± SD) was 25.7°C ± 4.1 ranging between 24.1°C and 26.6°C across years. The greatest instantaneous rate of change in Diptera biomass occurred at 18.3°C (Fig. 3, A and B). Around this critical threshold, a two degree decrease or increase in maximum daily temperature led to a 13.6% drop or 15.0% rise in Diptera biomass, respectively. Based on this threshold, 8.2% of farm visits followed at least one cold snap (Appendix S2: Fig. S2). A second, less steep maximum of the first derivative was noted at 29.2°C resulting in a marked decrease in Diptera biomass (Appendix S1: Fig. S2). We further observed that the daily likelihood to observe a cold snap has increased slightly over the course of this data set (Appendix S4: Table S1 and Fig. S1), and that they were more likely to occur in less agro-intensive landscapes (Appendix S4: Table S2 and Fig. S2).

**Fig. 3:**
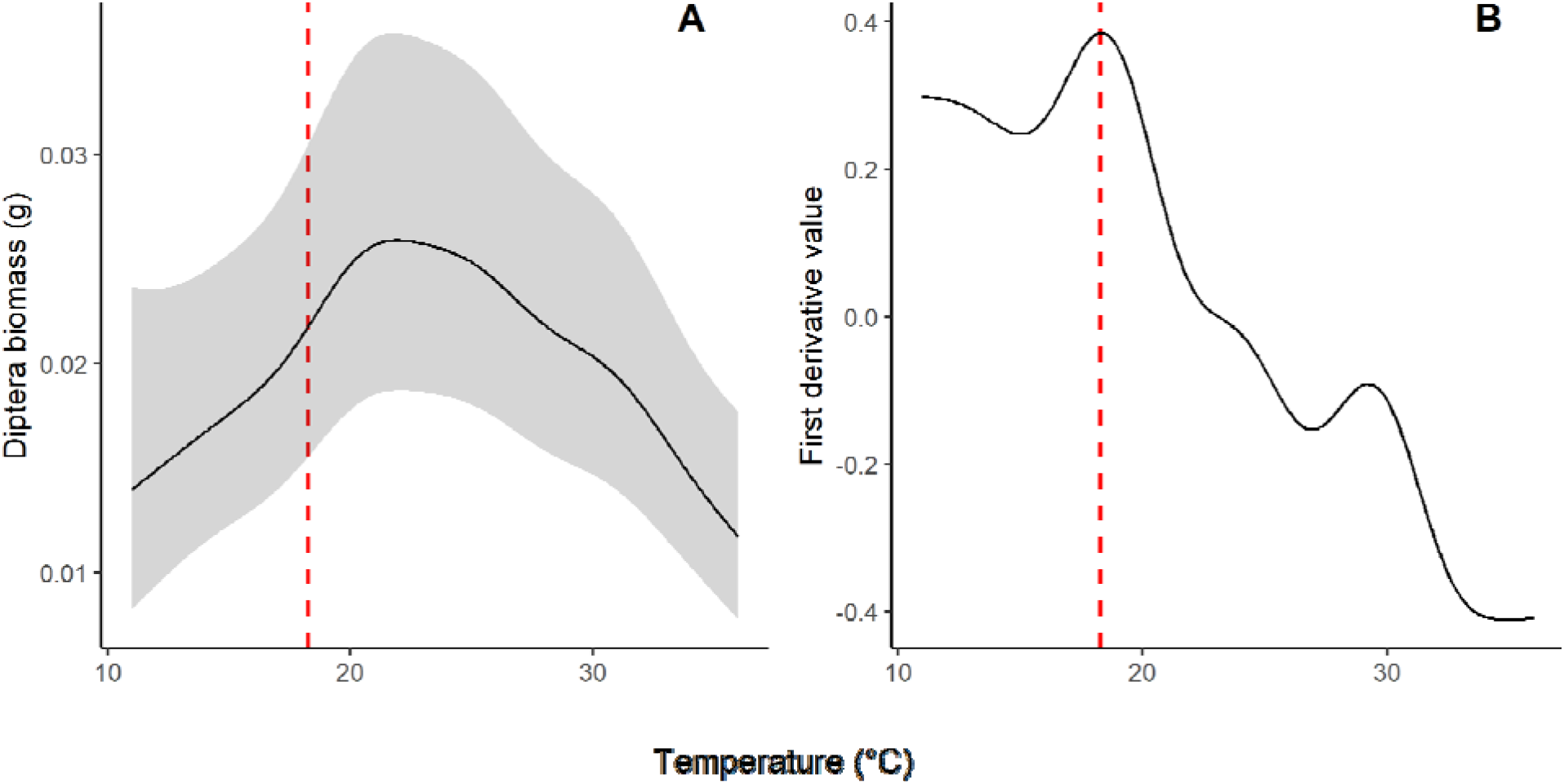
Functional relationship between Diptera biomass and the mean maximum temperature between the day of and day prior to sample collection (A) and the first derivative values of this relationship (B). Diptera biomass was derived with GAMMs using a Gamma distribution and a log link function. Vertical dashed line indicates the temperature at which the maximum of the first derivative was observed. Model controlled for, as tensor product smoothed terms, the Julian date and the interaction between sites score values (Comp.1 * Comp.2), and the total precipitation two days prior to insect collection (N=15,916 samples). Model also included year and farm IDs as random effects.

### Effects of inclement weather on fledging success

We monitored the breeding activity of 1,897 broods across the 40 farms and 11 breeding seasons. Overall mean (± SD) fledging success (i.e., ratio of the brood that fledged) was 0.74 ± 0.38, varying between 0.63 and 0.88 among years. We observed that the average hatching date advanced by ~1.1 days over the study period (Appendix S4: Table S3 and Fig. S3). Overall mean number of cold snap days (± SD) experienced by a brood during its first 12 days was 0.9 ± 1.1 days and varied between 0.2 and 2.3 days among years (Fig. 4). At least 52% of broods experienced at least one cold snap day throughout the duration of the study; 25%, 11%, and 5% of broods experienced at least 2, 3, and 4 cold snap days, respectively. Mean daily precipitation during the first 12 days (± SD) was 7.4 ± 4.9 ml, ranging between 2.4 and 13.6 ml across years. The correlation between the number of cold snap days and mean daily precipitation experienced by a brood was relatively low (*r*=0.18). For a more detailed look at the appearance of cold snaps and precipitation along the agricultural gradient of this data set see Appendix S2 (Fig. S3).

**Fig. 4:**
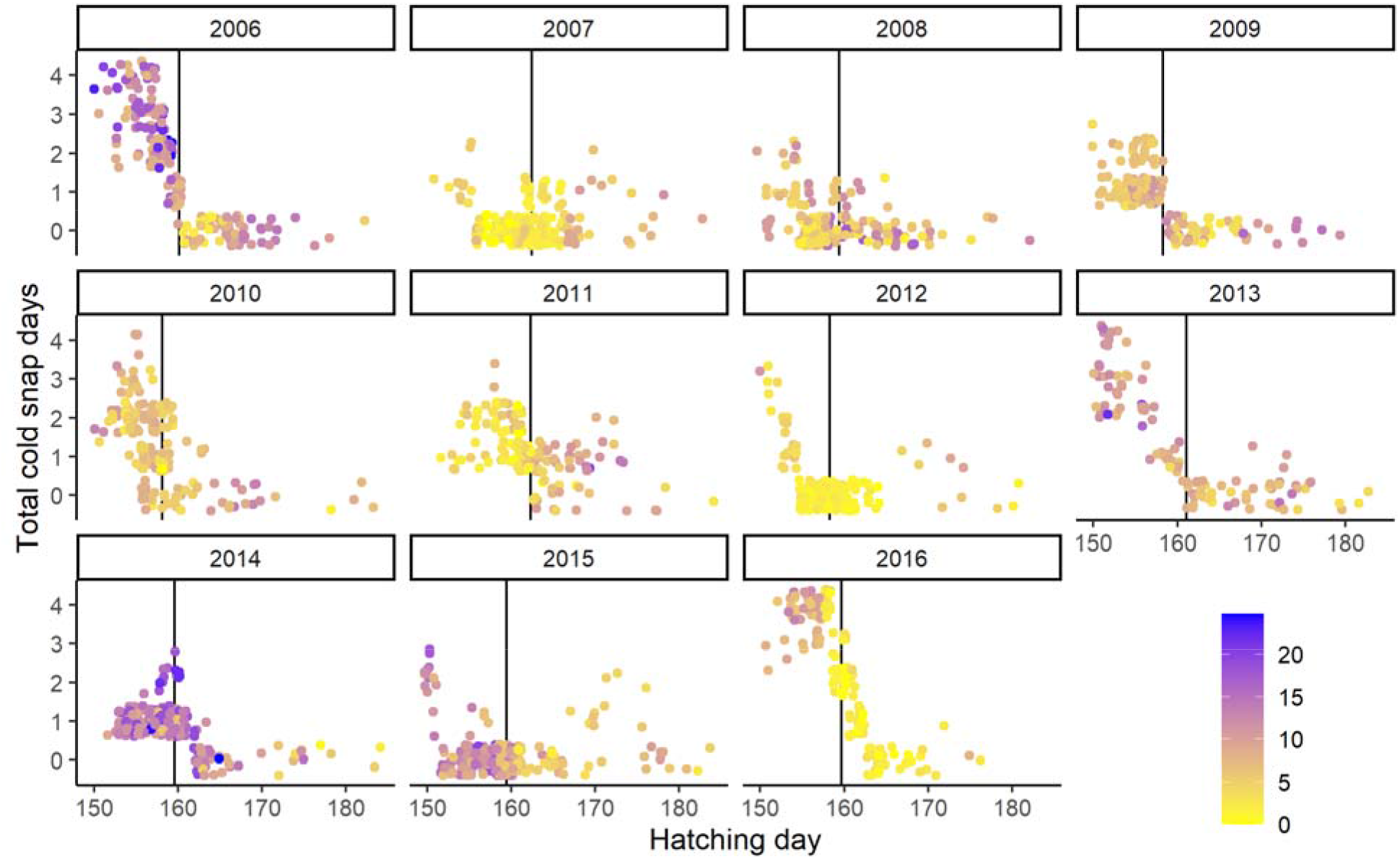
Number of cold snap days (jittered) experienced by Tree Swallow broods (N=1,897) according to their hatching dates (1 June and 15 July are Julian 152 and 196, respectively) between 2006 and 2016. Cold snap days were days on which maximum temperature on the farm did not surpass 18.3°C. Point color represents the mean daily precipitation volumes (ml) that broods experienced over a 12-day period after hatching. Black vertical line in the background is the mean hatching date of that breeding season.

Models incorporating an interaction between Comp.1 and the number of cold snap days (combined AICc *w* > 0.99) were substantially supported over Base (ΔAICc > 16.15, Appendix S2: Table S1). Fledging success decreased with the number of cold snap days (Fig. 5). Fledging success was on average 36.2% lower when broods experienced four cold snap days and average precipitation in their first 12 days instead of none. Increasing precipitation had a negative effect on fledging success, especially when broods were reared during windows containing both several cold snap days and high average daily amounts of precipitation (Appendix S2: Fig. S4). Broods subjected to four cold snap days versus none showed up to a 46.0% reduction in fledging success if exposed to the 95^th^ percentile of observed values for mean daily precipitation. The effects of cold snap days varied with landscape context (Fig. 5). For broods experiencing increased cold snap days, higher levels of agro-intensive cover was associated with further reductions in fledging success. When broods incurred the average number of cold snap days and mean daily precipitation there was no observable difference in fledging success along the agricultural gradient. However, this pattern changed to fledging success being 43.6% lower in agro-intensive landscapes when broods experienced four cold snap days and average precipitation (Appendix S2: Fig. S5). The synergistic negative effect of the number of cold snap days and high amount of precipitation on fledging success was, however, far greater in agro-intensive landscapes than in forage landscapes. In the absence of precipitation, the negative effect of increasing cold snap days was substantial only in agro-intensive landscapes, but similarly detrimental over the entire intensification gradient under high precipitation levels (Fig. 6). Lastly, fledging success increased with local prey availability and percent cover of surface water within 3 km, it decreased with increasing hatching day and brood size, and was lower for SY than ASY females (Fig. 5).

**Fig. 5:**
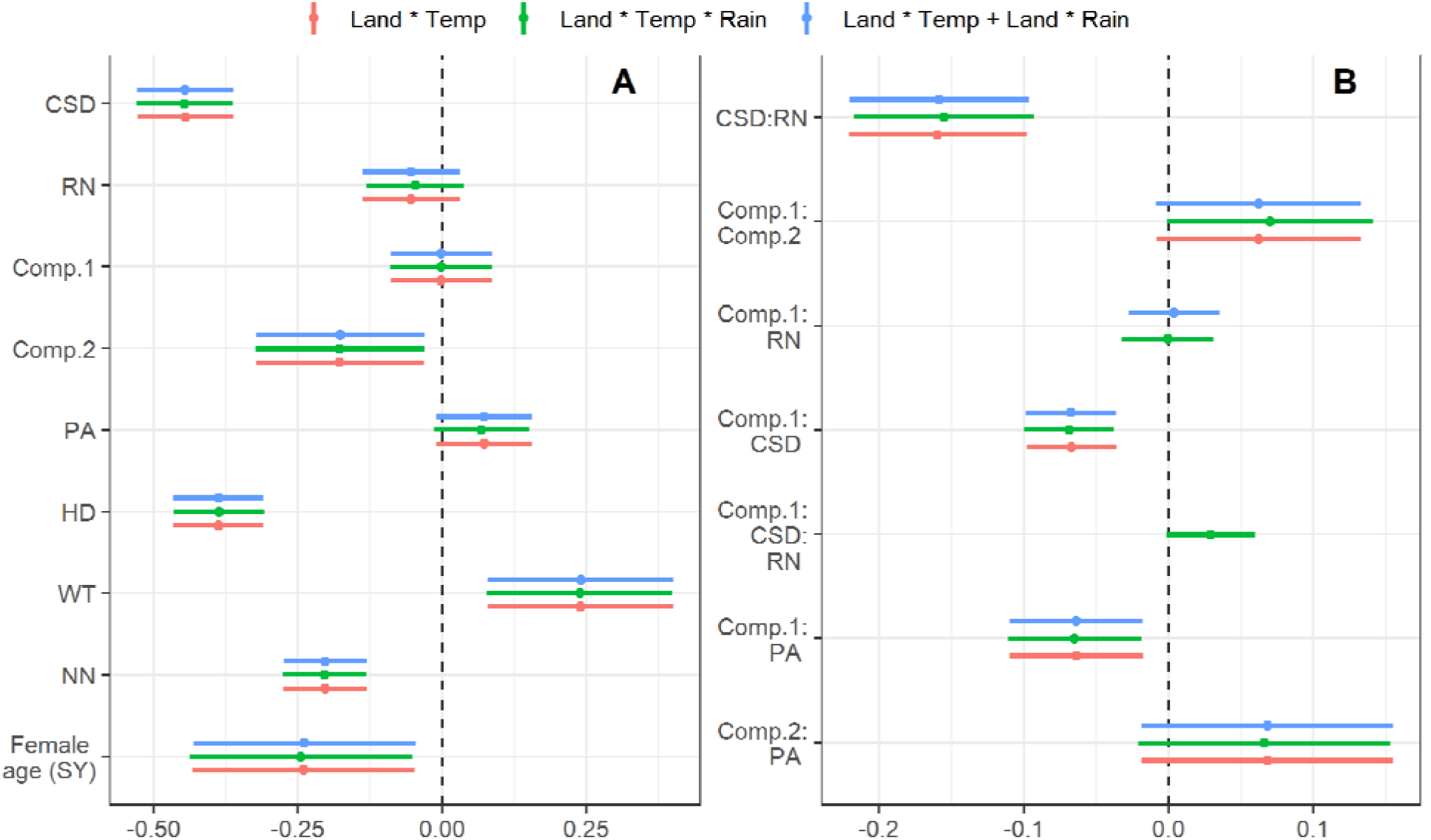
Standardized coefficient estimates and 95% confidence intervals of the three top ranking models, with combined *w* > 0.99 and max ΔAICc = 1.2. Models quantified the effect of covariates on the fledging success of Tree Swallow broods (N=1,897) raised across a gradient of agricultural intensification within southern Québec, Canada, between 2006 and 2016. See Table 1 for acronyms and summary statistics of fixed effects and Appendix S2: Table S1 for outcome of model selection. The GLMM with binomial response and logit link function included year, farm, and nest box IDs as random effects. Model weights of the top three supported models suggest they are comparable and thus are presented together.

**Fig. 6:**
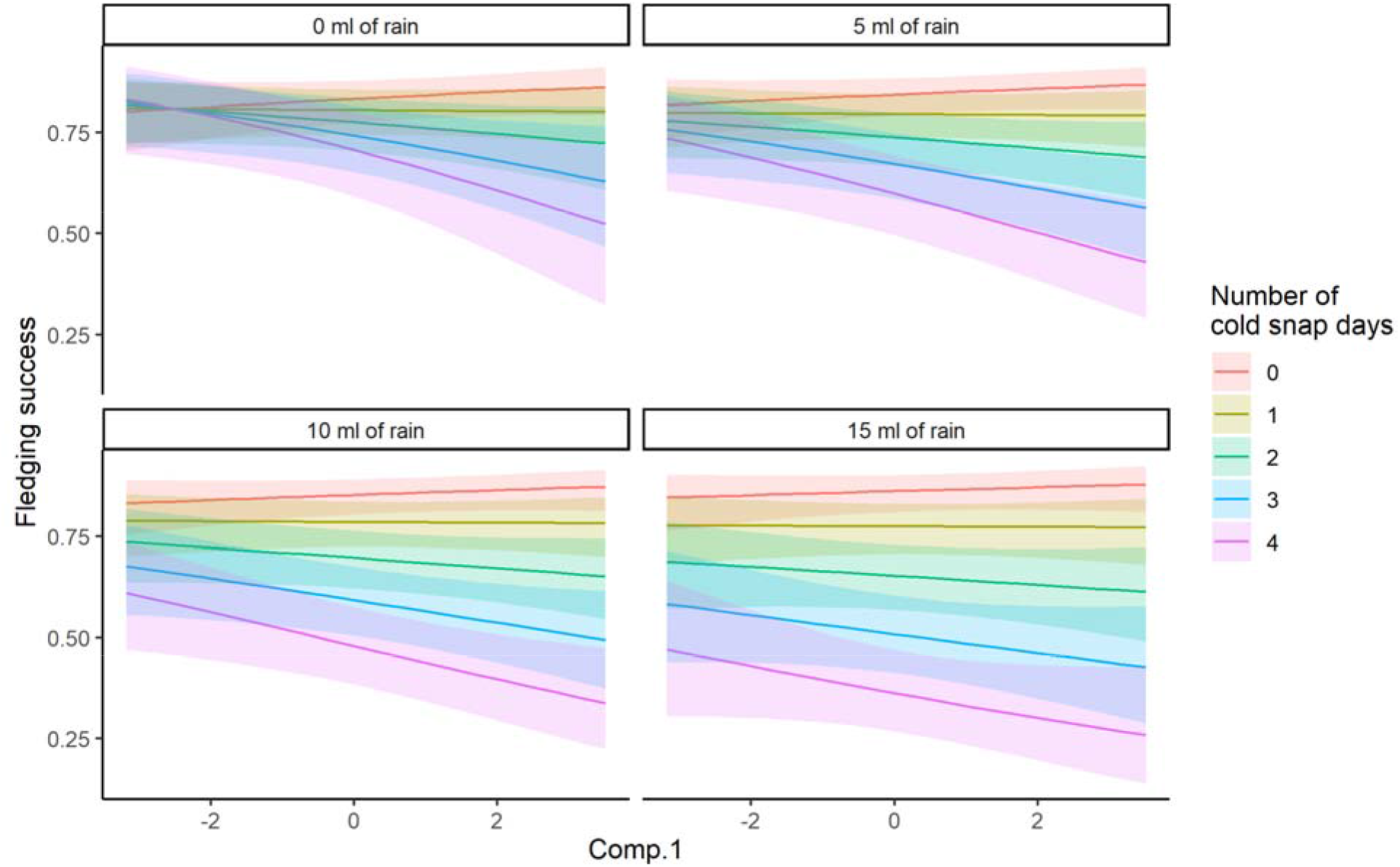
Unconditional model predictions (including unconditional 95% confidence intervals) of the effect of landscape context (Comp.1 in Fig. 2) on the fledging success of Tree Swallow broods (N=1,897) exposed to different numbers of cold snap days (days which maximum temperature < 18.3°C) and, in each panel, the mean daily precipitation volumes (ml) over 12 days post hatching. Predictions were made keeping all other covariates at their mean.

## Discussion

An outcome of climate change is the increased frequency and duration of weather events potentially reducing the availability of thermally sensitive prey resources (Rahmstorf and Coumou 2011, Wuebbles et al. 2014). These reductions, when occurring during critical life history stages (e.g., nestling periods), reduce the fitness of avian aerial insectivores (Pipoly et al. 2013, Moreno et al. 2015, Arbeiter et al. 2016). Though climate change influences a vast array of species and their interactions, another large-scale continued threat to biodiversity is anthropogenic reductions in habitat amounts and quality, notably through forestry and agriculture (Maxwell et al. 2016). We contended that the use of certain habitats may pose a risk to an animal’s ability to breed or survive through periods of reduced food availability and/or poor weather. We provide evidence that the negative impact of weather conditions expected from climate change on the annual breeding success of an aerial insectivore are exacerbated in habitats created by agricultural intensification.

The relationship between prey availability and mean maximum temperature, between the day of and day prior to sample collection, revealed the rate of change in prey availability was maximized at a temperature (18.3°C) similar to that deemed critical in past works (18.5°C; Winkler et al. 2013, Cox et al. 2019, Shipley et al. 2020). Such similarity may highlight a generality to temperatures impactful to Tree Swallows and their prey. We further found that this temperature threshold did not vary across the large-scale gradient of agricultural intensification studied here.

In accordance with previous studies, the appearance and frequency of cold snaps during the 12-day post hatching period of each brood led to reduced fledging success of Tree Swallows. This effect increased when multiple cold snaps and increased rain occurred during the 12-day post-hatching period. However, the overall negative effect of experiencing increasing precipitation appeared to be much weaker than increasing number of cold snaps. The impact of cold snaps likely acts on fledging success both through variation in resource availability, and by increased thermoregulatory demand (Sauve et al. 2021). While precipitation, because nests are protected within nest boxes, likely only impacts nestling survival through the former mechanism. This result is further evidenced by the role of the agricultural gradient in this relationship. Most notably, the negative effect of poor weather was stronger for breeding attempts in more agro-intensive landscapes. Such a result suggests features associated to these landscapes exacerbate the consequences of broods experiencing poor weather.

It should be mentioned that under very dry and warm conditions, fledging success appears to be slightly better in the most agro-intensive landscapes (7 % increase along the Comp.1 axis in Fig. 1 with 0 ml of rain). This relationship may highlight that agricultural landcover is only influential to fledging success during certain weather conditions as highlighted by the significant support of models incorporating interaction terms representing such a relationship. There is no clear explanation for this increase in fledging success in agro-intensive areas under these conditions. However, one potential hypothesis is that hay harvesting, which causes insect mortality and which timing depends on weather conditions, may affect the differences in quality between agricultural habitats (Humbert et al. 2009). However, food provisioning rates appear to be the lowest within the most agro-intensive parts of this study system (Garrett et al. 2021b). If the lowered food provisioning in these conditions are indicative of greater costs for food provisioning parents, then these costs may carry over and influence the survival and lifetime reproductive success of these parents. Therefore, under optimal weather conditions, fledging success may be slightly greater in more agro-intensive landscapes, and yet negatively impact the survival of adults when compared to forage landscapes. It should be stated that in our data set, broods experiencing 0 cold snap days and mean precipitation predominantly occurred within more agro-intensive landscapes, and that the daily likelihood to experience a cold snap was greatest in more forage landscapes. This is likely due to the elevational change occurring along the agricultural gradient. Taken together this may indicate that we underestimate the consequences of rearing young within agro-intensive landscapes, as environmental conditions from this data set appeared spatially worse for those broods in forage focused landscapes.

We propose two non-mutually exclusive mechanisms for the finding that broods experiencing inclement weather succumbed to greater mortality in more agro-intensive landscapes compared to forage focused landscapes: the structural characteristics of agro-intensive landscapes and the likelihood of exposure to agrochemical agents. Successfully acquiring food during periods of poor weather potentially result in greater costs for food provisioning parents, inasmuch as the increased foraging effort needed to overcome the reduced availability of prey likely results in reduced body condition (Hinsley 2000, Evens et al. 2018). Parents may possibly forgo foraging during poor weather periods in order to brood nestlings and provide food at greater rates once poor weather has subsided (Cox et al. 2019). Given the homogeneity of intensively-managed agricultural landscapes, prey patches may be fewer or farther apart therein (Benton et al. 2003, Fahrig et al. 2011). Furthermore, agro-intensive landscapes tend to be denuded of structural habitats [e.g., woodlots and hedgerows; Fig. 1; (Benton et al. 2003, Fahrig et al. 2011)]. In addition to being prey habitat, these features may block wind, and their removal may limit prey aggregation (Evans et al. 2003, Grüebler et al. 2008) and facilitate heat dissipation of nests and adults (Heenan 2013, Tapper et al. 2020a). Therefore, the impacts of experiencing periods of inclement weather in forage dominated landscapes may be less harsh than in more agro-intensive landscapes.

Evidence suggests that agrochemical agents such as herbicides (e.g., glyphosate, atrazine, S-metolachlor, etc.), neonicotinoid insecticides (e.g., clothianidin, thiacloprid, thiametoxam, etc.), and fertilizers can alter avian physiology in ways that potentially hinder an individual’s capacity to respond to poor weather (Mayne et al. 2005, Mineau and Palmer 2013, Gibbons et al. 2014, Lopez-Antia et al. 2015), most notably through a reduction in thermoregulatory capacity (Grue et al. 1997). These same agents may also influence the ability for food provisioning parents to optimize foraging strategies by inducing anorexia and impairing both locomotor and navigational function (Eng et al. 2019). These factors illustrate that within more agro-intensive landscapes, food provisioning parents may present longer foraging bouts that potentially come at a cost to their own body condition, while also reducing the time spent brooding nestlings (Garrett et al. 2021b).

Our conclusions, as well as predicted impacts of climate change on aerial insectivores, assume food provisioning adults are unable or less capable of foraging during periods of poor weather. However, fine scale movements of small passerines are becoming less difficult to observe (Elgin et al. 2020, Garrett et al. 2021b). Understanding such movements may further elucidate the combined roles of breeding habitat quality and predictions of climate change on animal behavior. For instance, the increased food provisioning of nestlings by adults, following periods of poor weather (Cox et al. 2019), may be facilitated by foraging in more profitable habitat patches (Elgin et al. 2020, Geary et al. 2020, Garrett et al. 2021b). In the context of climate change, the combined roles of poor weather and landscape structure on foraging responses is still unknown. Validation of these assumptions will be imperative as the effectiveness of food provisioning, during periods of poor weather, likely varies across a gradient of breeding landscape quality. Finally, a focus here was on the effects of cold snaps, yet another growing concern of climate change are periods of high ambient temperature [i.e., heat waves; (Perkins-Kirkpatrick and Lewis 2020)]. The energetic expenditure of food provisioning Tree Swallows is potentially constrained by an upper thermal limit (Tapper et al. 2020b). Moreover, prey availability decreased sharply above 30°C (Appendix S1: Fig. S2) in our study system. Food provisioning parents may thus be hindered and the effects of breeding within more agro-intensive landscapes may further be exacerbated during heat waves. Future work should focus on evaluating this hypothesis, as reduced food provisioning potentially results in reduced nestling growth and survival (Cox et al. 2019, Garrett et al. 2021b).

Finally, our focus on the relationship between a threshold temperature and nestling survival utilized a single temperature and assumed that experiencing days falling below this value impacts both the amount of food resources coming into nest boxes and the thermoregulatory capacity of nestlings. However, our threshold temperature of 18.3°C occurs multiple times during a breeding season and thus selective processes would likely favor more robust capacity of nestlings to survive such temperatures. Yet, we found strong support that nestling mortality increased when broods experienced increasing number of cold snaps. We propose that there is likely two threshold temperatures (Appendix S3: Fig. S1), one influential to the availability of prey resources and another influential to nestling’s thermoregulator capacity (Stodola et al. 2010, Pérez et al. 2016, de Zwaan et al. 2019), as proposed by Winkler et al (2013). Future work needs to explicitly evaluate this hypothesis across similar gradients of agricultural intensification.

## Conclusion and applications

Aerial insectivore populations are declining in several parts of their North American and European breeding grounds (Spiller and Dettmers 2019, Rosenberg et al. 2019, Bowler et al. 2019). Here we provided evidence that two of the primary hypotheses explaining these reductions, especially for aerial insectivores, may interact to further intensify declines. We propose that future work investigating insectivorous bird population declines should explicitly focus not only on climate change or anthropogenic reductions or alterations of breeding habitats, but also on how these factors may interact with one another. Furthermore, understanding the various mechanisms that lead to increased nestling mortality during inclement weather or within more agro-intensive landscapes will be critical as they potentially identify drivers under anthropogenic control.

## Acknowledgments

We are indebted to the farm owners who kindly accepted to partake in our long-term study since 2004. Sincere thanks to the many graduate students, field and lab assistants who helped collect, enter, and proof data throughout the years, notably with respect to insect sample processing. This work was conducted under the approval of the animal care committee of the Université de Sherbrooke and was financially supported by Natural Sciences and Engineering Research Council of Canada (NSERC) discovery grants to FP, DG and MB, two team research grants from the Fonds de recherche du Québec—Nature et technologies (FRQNT) to FP, DG and MB, by the Canada Research Chairs program to FP and MB, as well as by New Opportunities Funds of the Canadian Foundation for Innovation (FCI) to FP, DG and MB, the Canadian Wildlife Service of Environment and Climate Change Canada and the Université de Sherbrooke.

