## Appendix S1 for "Interacting effects of cold snaps, rain, and agriculture on the fledging success of a declining aerial insectivore"

#### Appendix S1: Defining cold snaps

#### Characterization of cold snap temperatures

We desired cold snap values to be grounded in the functional response between local prey availability and temperature. We used generalized additive mixed effects models (GAMMs) and the bam() function from the mgcv package (Wood 2015) in R to complete this modeling endeavor. We first modeled the relationship between Diptera biomass and temperature, as insects require threshold temperature values to be met before they are active (Williams 1961). We then determined the temperature at which the instantaneous rate of Diptera biomass change was greatest. Peak instantaneous rate was estimated as the maximum of the first derivative of the relationship between Diptera biomass and temperature. We chose peak rate as temperatures around this critical threshold likely result in dramatic differences in prey availability. Diptera biomass demonstrated right skewness and models incorporated a Gamma distribution with a log link function. In all subsequent models, we include the Julian date of sample collection as a tensor product smooth (k=10) to account for phenological variations in biomass as well as precipitation. We further included both the year and farm IDs as random effects (s (x, bs=’re’)) in order to control for the hierarchical structure of our sampling design. In order to evaluate overall trends, we kept the basis dimension of temperature low (k=10).

We evaluated if critical temperatures, represented as the mean daily maximum temperature between the day of and day prior to sample collection, were landscape dependent, and thus explored three models including site score values and temperature. In all models, site scores were treated as a bivariate smooth using a tensor product smooth (s (Comp.1, Comp.2, k=100)). We hypothesized that the effect of temperature on Diptera biomass was independent of the landscape context, and thus included the temperature term plus site scores (Temp + Land). We further hypothesized that the effect of temperature on Diptera biomass varied according to a functional response specific to the landscape context (Temp * Land). We thus used a tensor-product interaction between landscape context and temperature values without an underlying functional response of temperature. This interaction effectively allows the functional response of temperature at each site score combination to not be constrained by any underlying pattern. Finally, we hypothesized that the effect of temperature on Diptera biomass varies with landscape context and yet follows an underlying trend (Temp + Temp * Land) and used a tensor-product interaction between landscape context and temperature. Including the smoothed term of temperature along with the three-way smooth effectively constrains the functional response between temperature and Diptera biomass to follow a similar trend with each combination of site score (Pedersen et al. 2019). Model selection indicated the model with the lowest ΔAIC (*w*>0.99) was a model (Temp + Temp * Land) including the three-way smoothed term between temperature and site scores, constraining the functional response of temperature to resemble an underlying shape (Table S1). Predictions from this model suggest the functional response between Diptera biomass and temperature varied only in magnitude and not in overall shape (Fig. S3). These minor deviations in the functional response had no bearing on the maximum of the first derivative across the agricultural intensification gradient. For example, the vertical black line in each panel of Fig. S3 indicates the maximum of the first derivative for each curve in the figure (representing different levels of site score; Comp.1 and Comp. 2 in Fig. 2). These results signify that the cold snap temperatures, derived by the maximum of the first derivative of the functional response between Diptera biomass and mean maximum daily temperature, remain the same across the agricultural gradient (i.e., 18.3 °C).

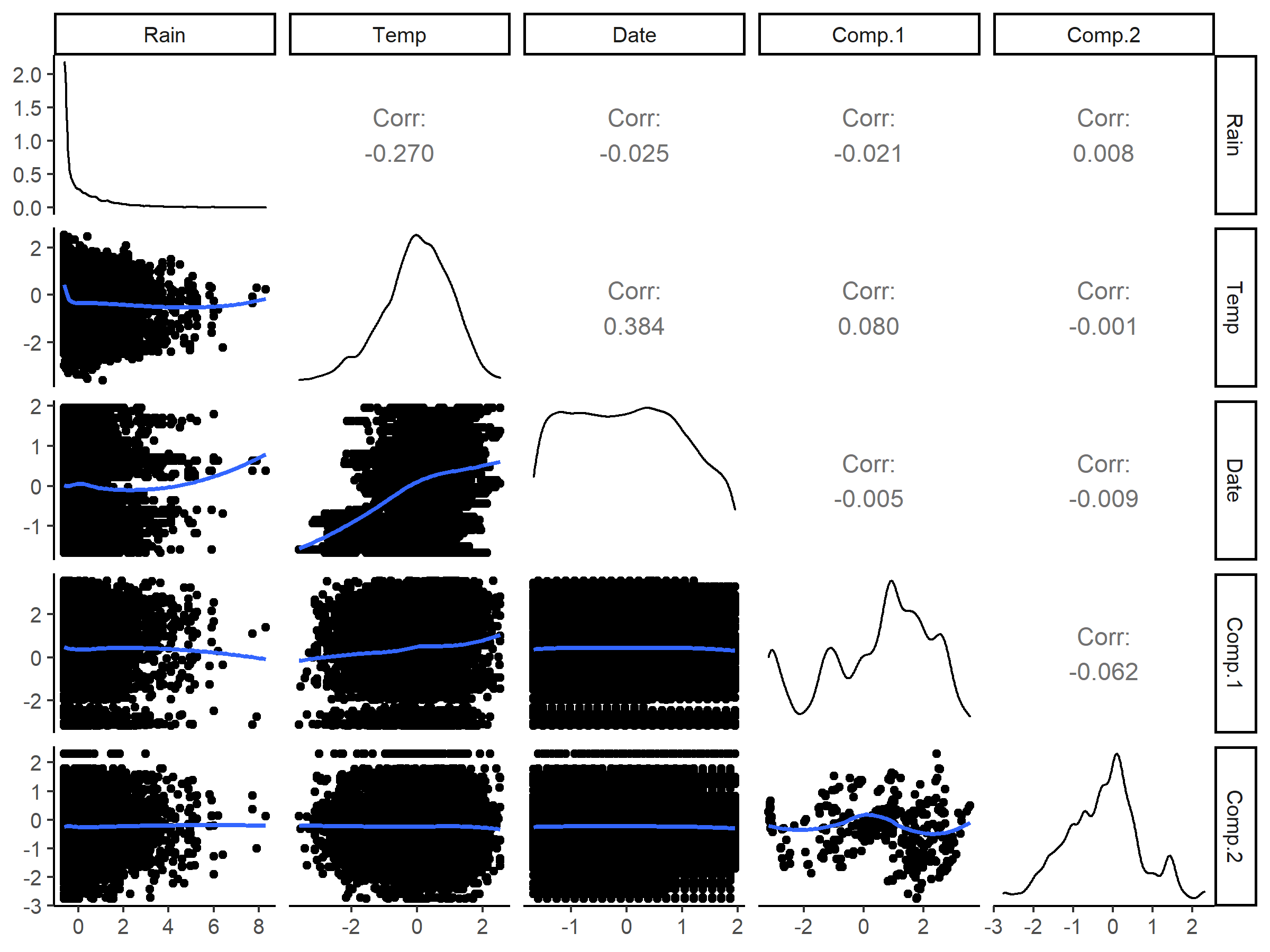

**Fig. S1:** Raw covariate data used in the modeling of Diptera biomass against weather across a gradient of agricultural intensification. Figure includes the pairwise correlations and functional relationships between each covariate. Blue smoothed curve in the lower left corner are predictions from a loess regression. **Table S1**: Outcome of model selection for the modeling of Diptera biomass against the mean maximum temperature between the day of and day prior to sample collection (TMax; N=15,497 samples) and landscape context (Comp.1 and Comp. 2 site scores; Fig. 2). Modeling involved generalized additive mixed effects models (GAMMs). Three right columns indicate the presence (+) of terms in the model. Confounding factors within each model include the Julian date of sample collection, as a tensor product smooth, and the year and farm IDs as random effects.

| Candidate models | ΔAICc | *w* | Deviance explained (%) | Comp.1* Comp.2 | Temp | Temp*  Comp.1*  Comp.2 |
| --- | --- | --- | --- | --- | --- | --- |
| Temp + Temp * Land | 0.00 | >0.99 | 19.14 | + | + | + |
| Temp + Land | 83.03 | 0 | 18.84 | + | + |  |
| Temp * Land | 934.77 | 0 | 16.19 |  |  | + |

**Table S2:** Concurvity measures of smoothed fixed effects terms in the top model. Concurvity is a value similar to collinearity but for non-linear models. Estimated values as calculated by Wood (2015) are shown. Values are bound between 0 (no concurvity) and 1 (worrying concurvity).

| Term | s(max1) | s(JJ) | s(Comp.1,Comp.2) | ti(max1,Comp.1,Comp.2) |
| --- | --- | --- | --- | --- |
| s(max1) |  | 0.12 | 0.00 | 0.00 |
| s(JJ) | 0.13 |  | 0.00 | 0.00 |
| s(Comp.1,Comp.2) | 0.11 | 0.02 |  | 0.01 |
| ti(max1,Comp.1,Comp.2) | 0.16 | 0.05 | 0.40 |  |

**Table S3:** Summary of top model predicting Diptera biomass as a function of the mean maximum daily temperature between the day of and the day prior to the collection of samples. Presented is the estimate, 95% CI, and p-value of the parametric terms and p-values for the relative effect of smoothed terms. Model used a Gamma distribution with a log link function on a data set (N = 15,916) occurring over a gradient of agricultural intensification represented by agricultural landcover around 40 farms over an 11-year period.

| Term | Estimate | CI | p-value |
| --- | --- | --- | --- |
| RN | -0.06 | (-0.08,-0.04) | **<0.001** |
| s(max1) |  |  | **<0.001** |
| s(JJ) |  |  | **<0.001** |
| s(Comp.1,Comp.2) |  |  | **<0.001** |
| ti(max1,Comp.1,Comp.2) |  |  | **0.006** |

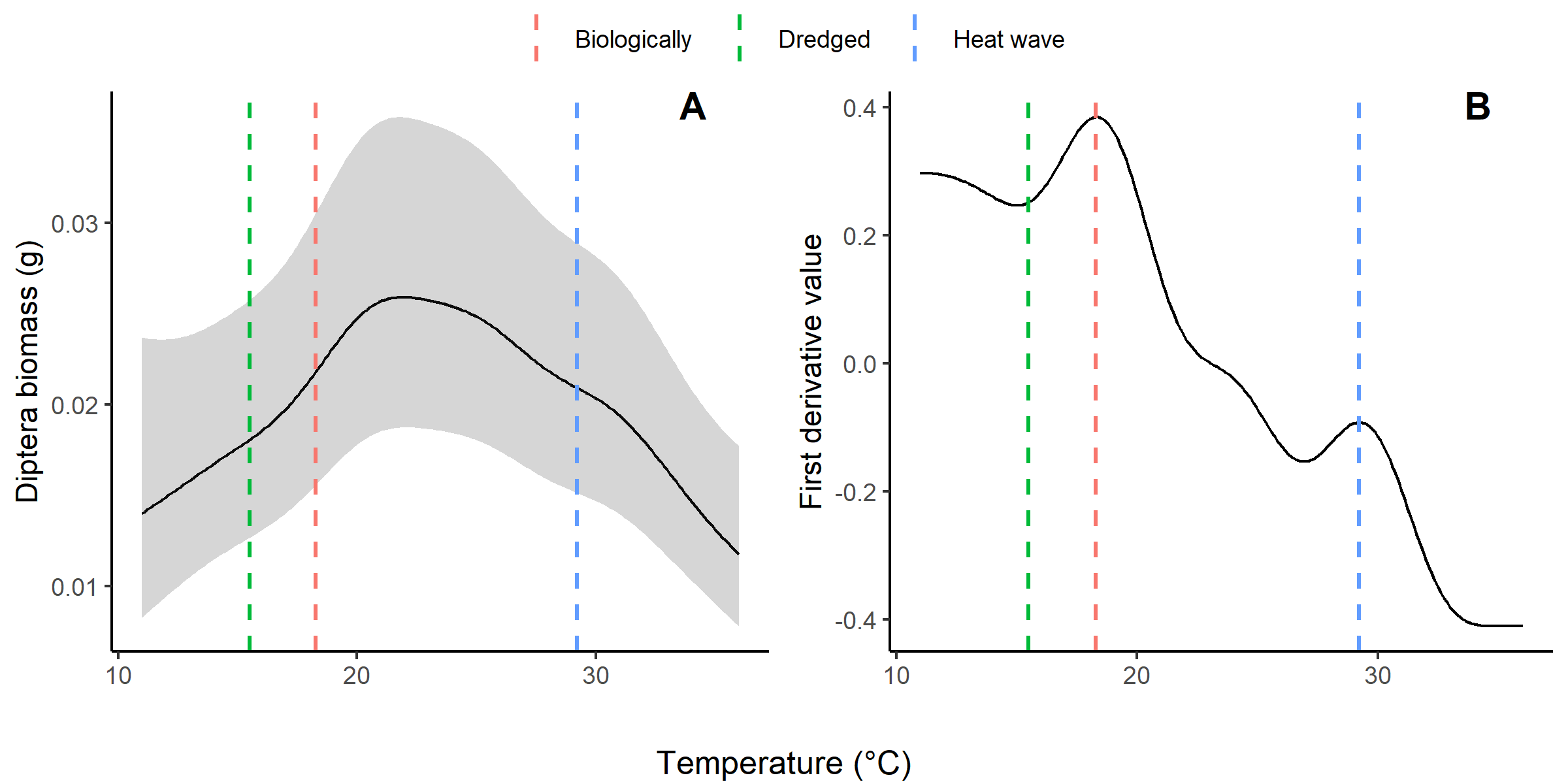

**Fig. S2**: The same as Fig. 2 showing the functional relationship between Diptera biomass and the mean maximum temperature between the day of and day prior to sample collection (A) and the first derivative values of this relationship (B). Diptera biomass was derived with GAMMs using a Gamma distribution and a log link function. Included are the three dashed lines representing the original cold snap temperature threshold from the text (red), the second peak in first derivative suggesting declines from heat (blue) and the data driven temperature from Appendix S3 (green).

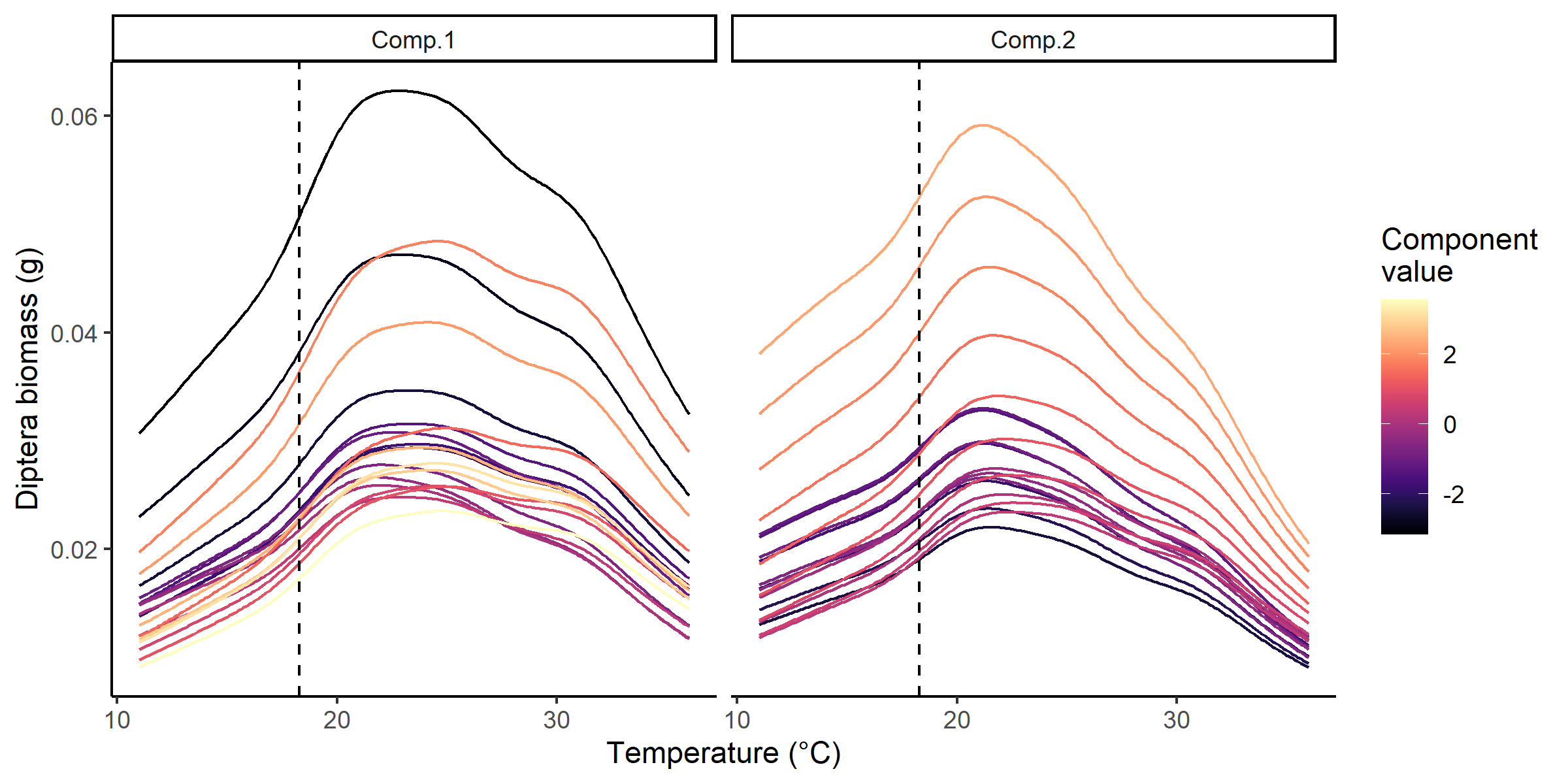

### Fig. S3: Model predictions from the most supported model (Temp + Temp * Land) of Diptera biomass against the mean maximum temperature between the day of and day prior to sample collection. Each curve represents the functional response between Diptera biomass and temperature for a different level of site score value (Comp.1 and Comp.2; Fig. 2). The underlying functional response remains similar across all site score combinations. Vertical black line of each panel indicates the temperature at which the maximum of the first derivative of the functional response between Diptera biomass and temperature was observed for each level of site score value.Supporting Information
