## Appendix S2 for "Interacting effects of cold snaps, rain, and agriculture on the fledging success of a declining aerial insectivore"

### Appendix S2: Supplemental tables and figures for fledging success

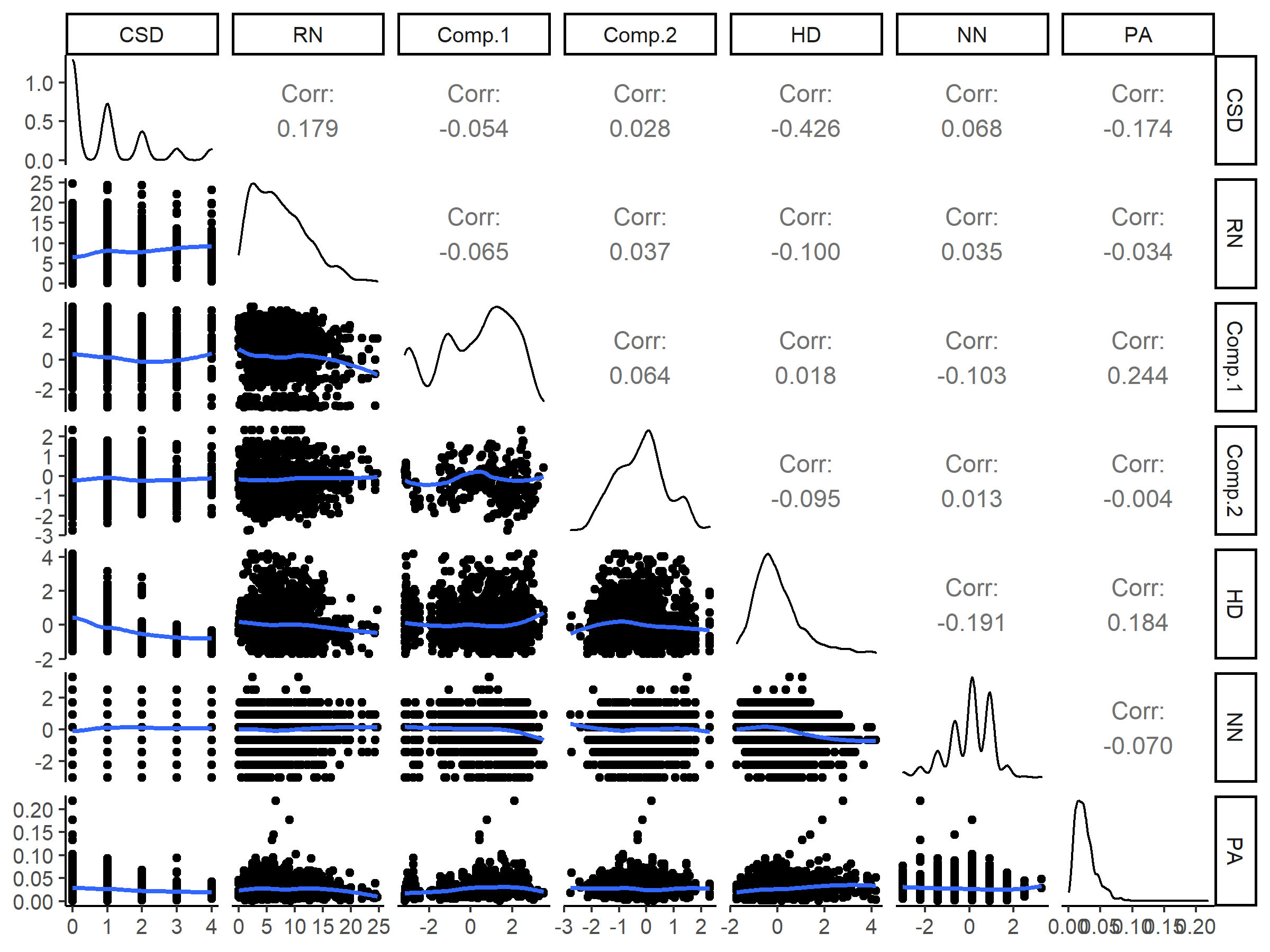

**Fig. S1:** Raw covariate data used in the analysis of fledging success across a gradient of agricultural intensification during inclement weather events. Figure includes the pairwise correlations and functional relationships between each covariate. Blue smoothed curve in the lower left corner are predictions from a loess regression.

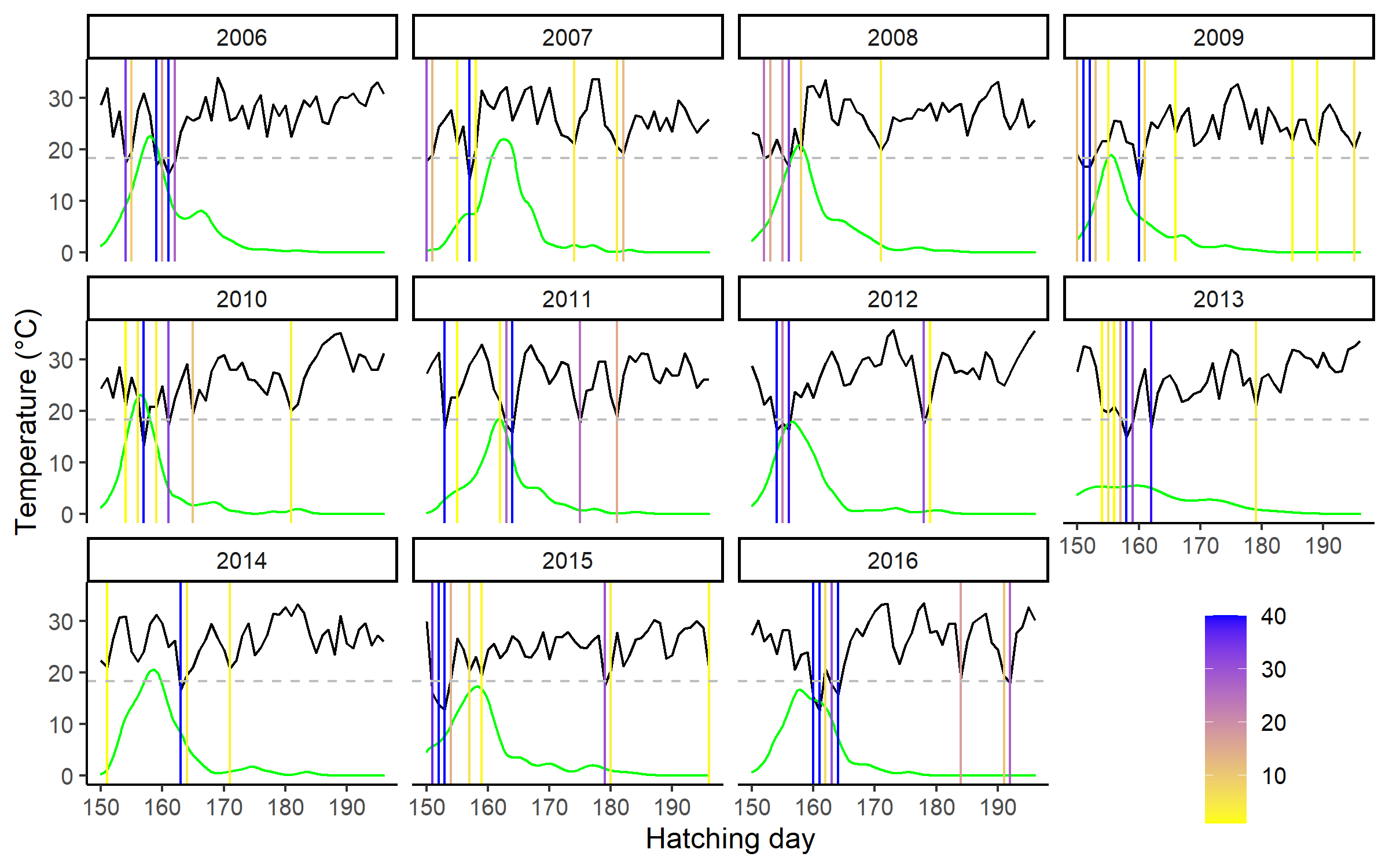

**Fig. S2**: Time series of mean maximum daily temperatures and hatching dates of Tree Swallow broods (N=1,897) between 2006 and 2016. Fitted black line in each panel is the mean maximum temperatures (calculated across all farms, N=40) each day. Horizonal dashed grey line is located at 18.3°C and designates the threshold temperature defining a cold snap. Green line delineates the frequency distribution of hatching dates. Vertical lines indicate the days on which at least one farm experienced a cold snap, and the line color specifies the number of farms subjected to a cold snap (range: 1-40 farms).

**Table S1**: Outcome of model selection regarding the fledging success of Tree swallow broods (N=1,897 broods) across a gradient of agricultural intensification within southern Québec, Canada, between 2006 and 2016. Fixed effects composing each variable group can be found in Table 1. Number of model parameters (K), difference in AICc values vs. the best model, and Akaike weight (*w*) are provided for each model composing the candidate set. Fledging success was modeled with GLMMs and binomial distribution with a log link function. Random effects included the Year (N=11), Farm (N=40), and Nest box ID (N=362) to account for hierarchical nature of the data.

| Candidate models | K | ΔAICc | *w* | Marginal R^2^ |
| --- | --- | --- | --- | --- |
| Land*Temp | 18 | 0.000 | 0.473 | 0.155 |
| Land*Temp*Rain | 20 | 0.596 | 0.351 | 0.157 |
| Land*Temp + Land*Rain | 19 | 1.983 | 0.176 | 0.155 |
| Base | 17 | 16.148 | 0.000 | 0.144 |
| Land*Temp*Rain | 18 | 18.111 | 0.000 | 0.144 |
| Null | 4 | 231.214 | 0.000 | 0.000 |

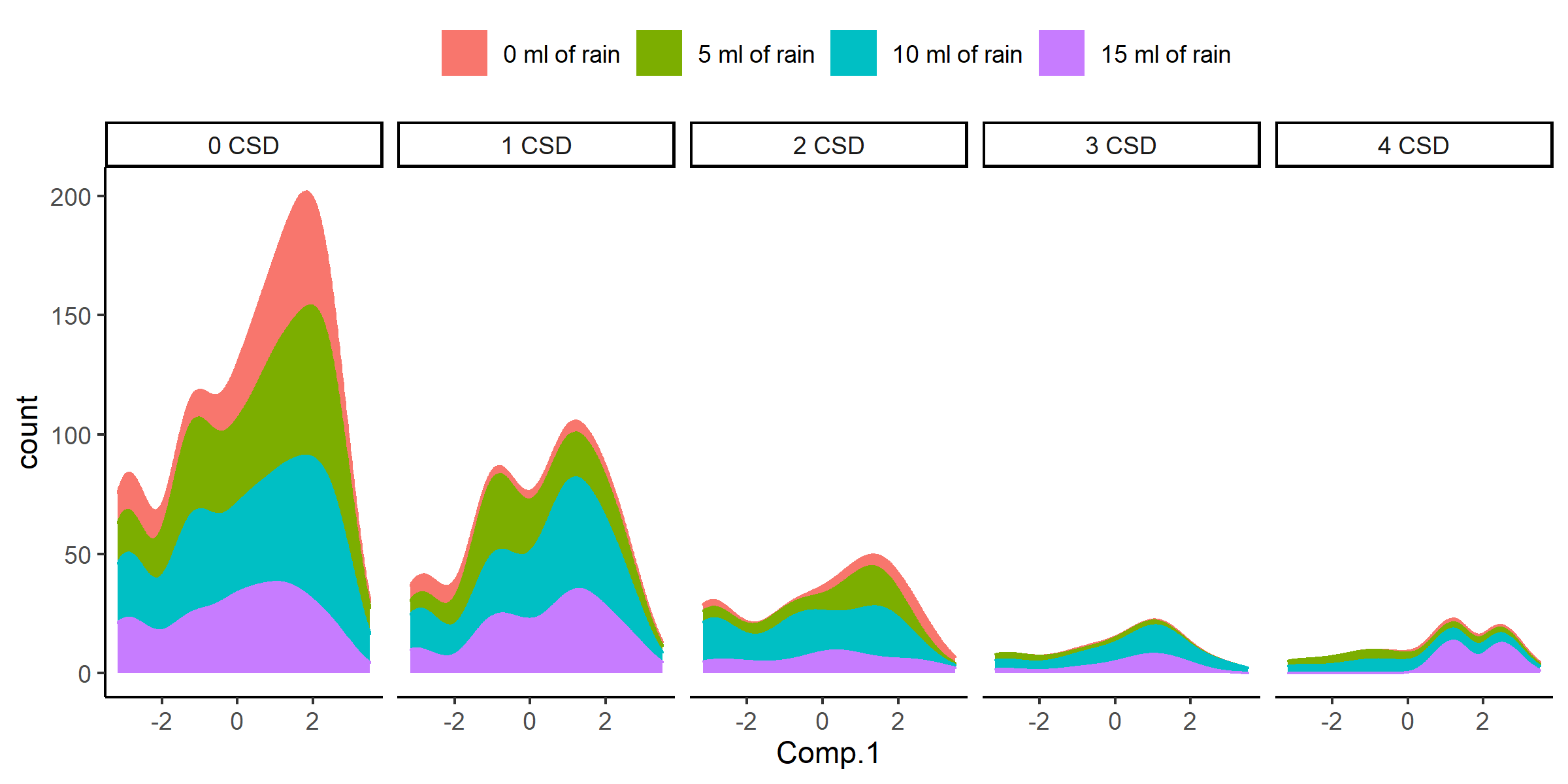

**Fig. S3:** Kernel density plots of the raw data on the occurrence of cold snap days and precipitation combinations. Colors are for observations binned by increasing mean precipitation by 5-ml increments (i.e., 0 ml are all the observations with mean of 0 ml during the 12-day post hatching period, while 5 ml are all the observations greater than 0 ml and less than 5 ml. etc.).

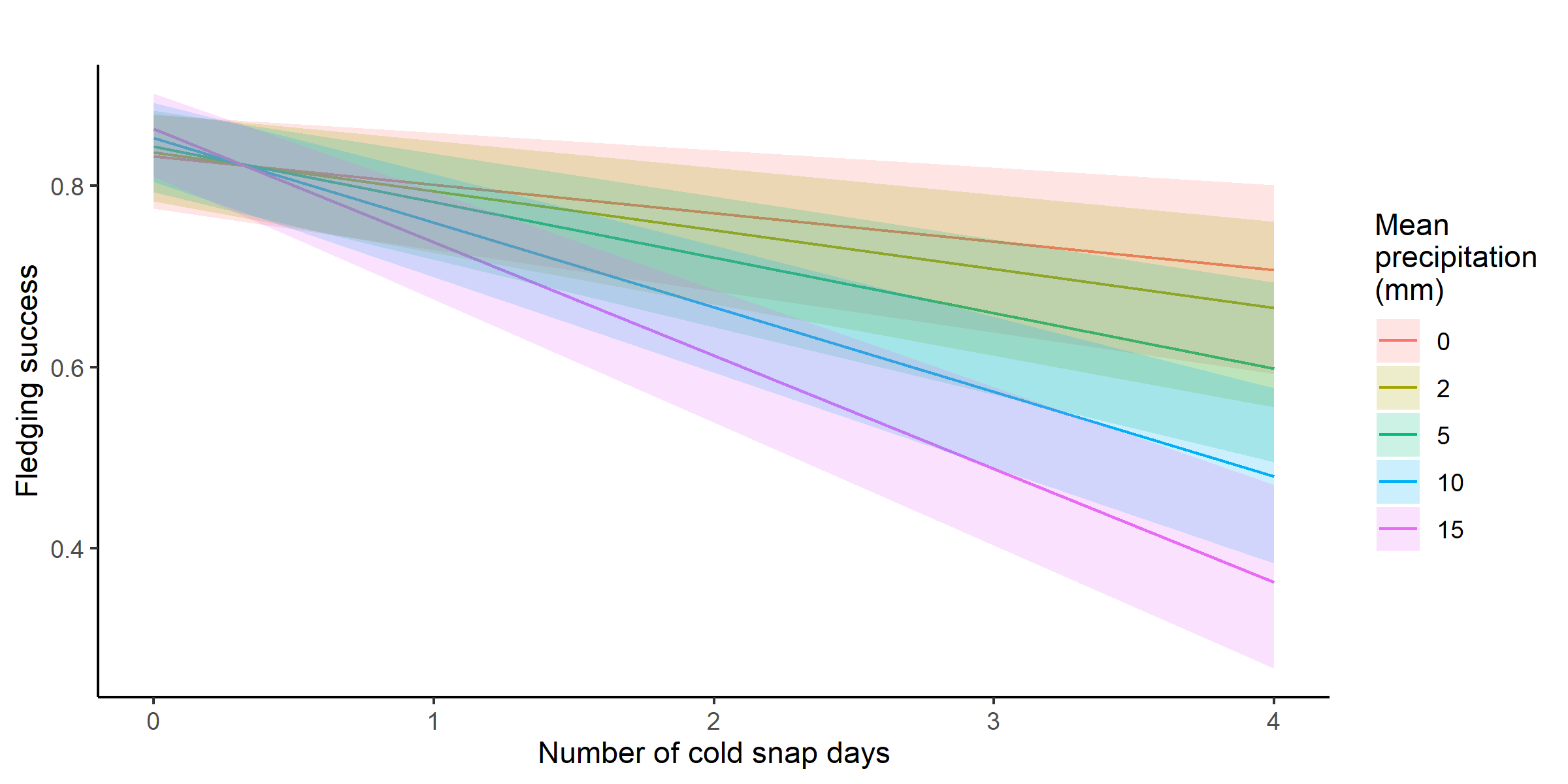

**Fig. S4**: Unconditional model predictions and associated 95% confidence intervals for the relationship between fledging success for broods experiencing increased number of cold snap days and precipitation during the 12-day post hatching period. All covariates, unless explicitly stated in the figure, were Z-transformed prior to analysis and set to their mean for the prediction.

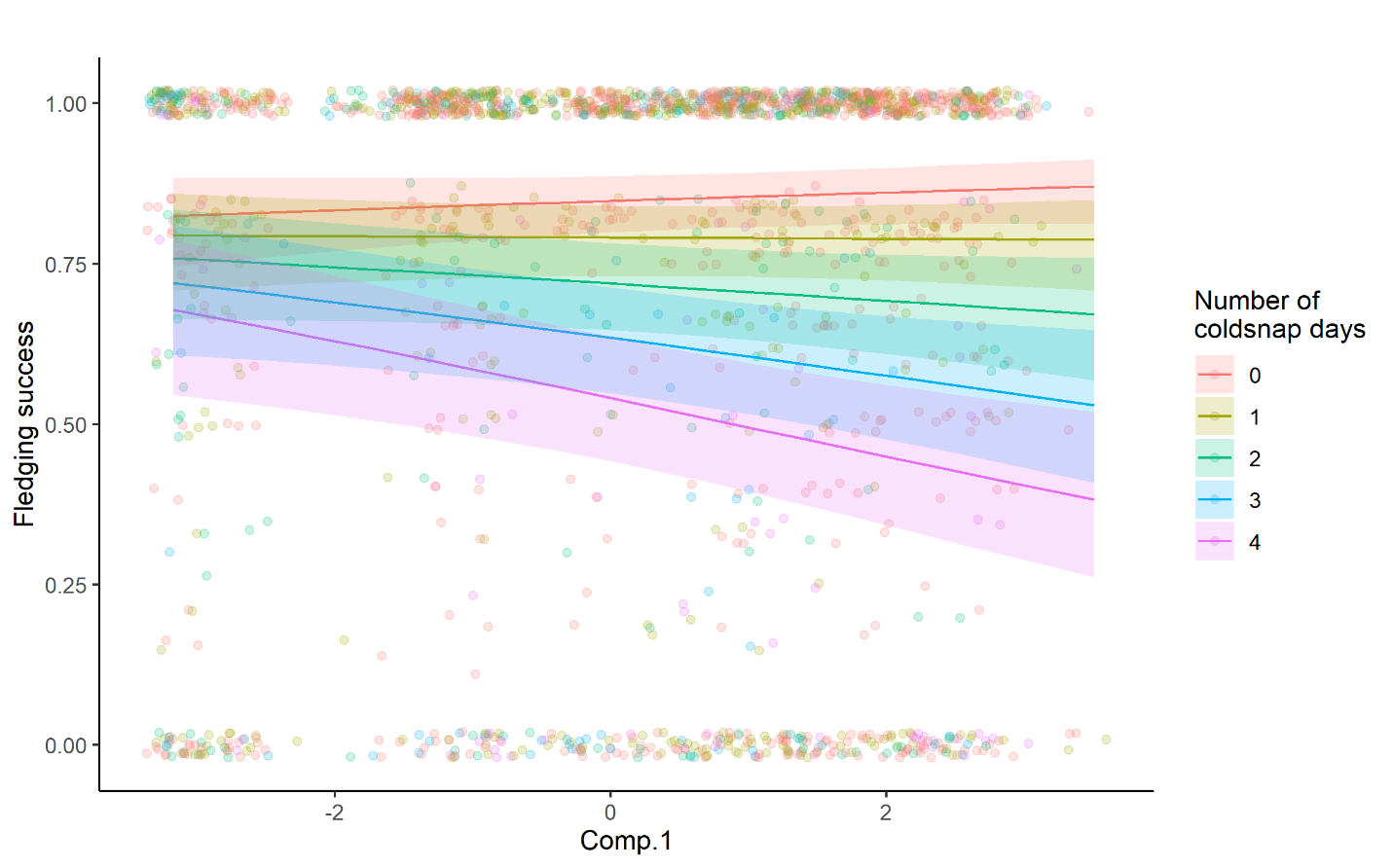

**Fig. S5:** Unconditional model predictions and associated 95% confidence intervals for the relationship between fledging success for broods experiencing increased number of cold snap days and mean precipitation during the 12-day post hatching period along the gradient of agricultural intensification. All covariates, unless explicitly stated in the figure, were Z-transformed prior to analysis and set to their mean for the prediction. Model predictions are plotted over the raw data.
