## Appendix S3 for "Interacting effects of cold snaps, rain, and agriculture on the fledging success of a declining aerial insectivore"

**Appendix S3: Supplemental analysis cold snap and precipitation definitions**

**Data driven threshold temperature and post-hatching window**

We desired weather variables to be grounded in conditions influential to the ecology of Tree Swallows to avoid spurious results and developing a hypothesis backed up by little physiological evidence (Burnham and Anderson 2002, Harrison et al. 2017). For these reasons, we chose a temperature influential to the physiology of insects and their availability (Taylor 1963, Roitberg and Mangel 2016), and a time window focussing on the period in which nestlings reached peak growth rate and body mass. These are both biologically sound assumptions (Burnham and Anderson 2002, Cade 2015). In this supplemental analysis, we underwent an intensive modeling endeavor to evaluate if our conclusions would hold up against other post-hatching windows and definitions of cold snaps. We compared a series of identical models (Land*Temp in Appendix S2 Table S1) in which only the time window and threshold temperature defining a cold snap varied. We compared all models with increasing post-hatching window between 4 and 16 days of age and a temperature threshold ranging between 13°C and 22°C. We then ranked all models by their AICc weight and compared these results to the AICc and threshold/window of our original model. We did find a second local minimum of *w* at a temperature threshold around 15.5°C and a post hatching window of 9 days (Fig. S1, point = “dredge”). This may indicate the possibility of two temperature thresholds, as hypothesised by Winkler et al. (2013). However, the appearance of this second minimum was arguably derived through data dredging (Harrison et al. 2017) and, though likely through hinderance of thermoregulation (de Zwaan et al. 2019, 2020, Glądalski et al. 2020), this temperature currently has little physiological support for Tree Swallows. It should also be highlighted that all coefficient estimates from the data driven top model are the same in overall magnitude and direction from the original model including both the non-interaction terms (Fig. S2) and interaction terms (Fig. S3). Therefore, interpretation of overall effects and predictions from this model would lead to similar conclusions derived based on the biological relationships between Tree Swallows and their prey we had previously used.

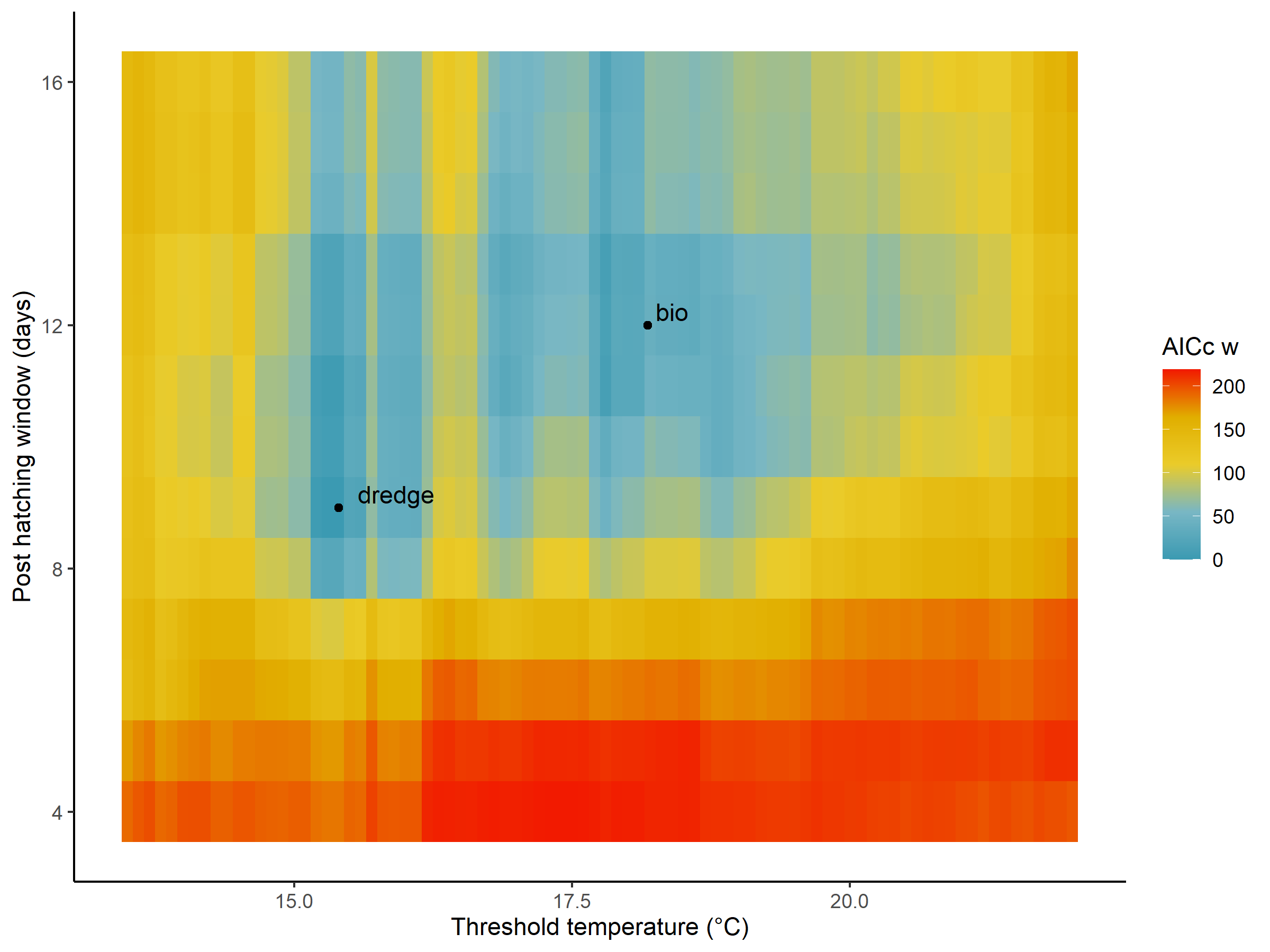

**Fig. S1**: Post hoc dredging analysis assessing the optimal threshold temperature determining fledging success. Presented is the ΔAICc of models given combinations of post hatching window and threshold temperature. The two points represent: the combination used in the text body (bio) and the combination of the model with the lowest AICc (dredge).

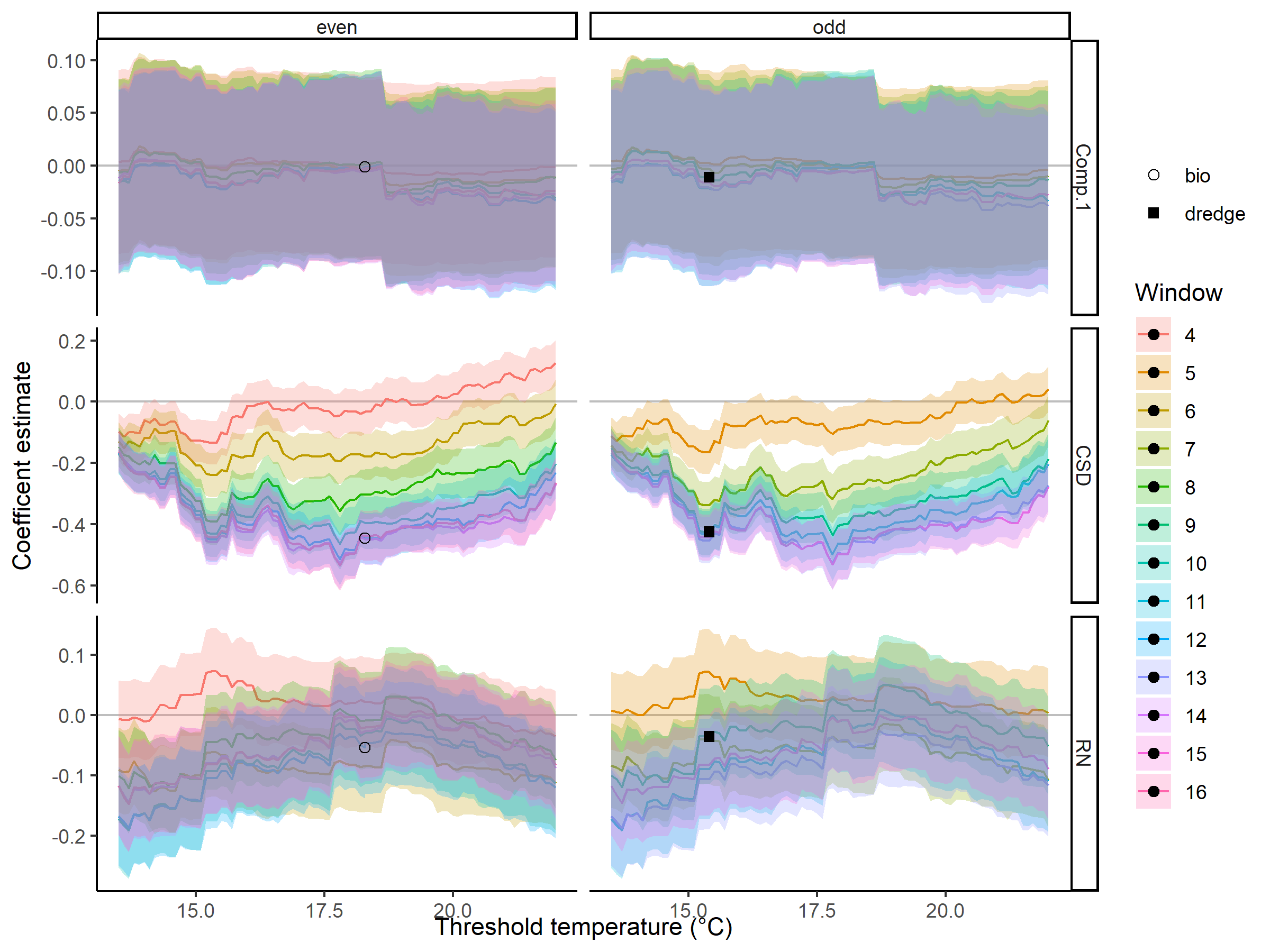

**Fig. S2**: Post hoc dredging analysis assessing the optimal threshold temperature determining fledging success. Presented are the coefficient estimates and 95% CI for the main non-interaction terms of interest with threshold temperatures increasing by 0.1°C. Presented colors are the post-hatching window increasing between 4 and 16 days post hatching. The two points represent: the combination used in the text body (bio) and the combination of the model with the lowest AICc (dredge). The two panels are the results separated by if the post hatching window was either an odd or evenly numbered window.

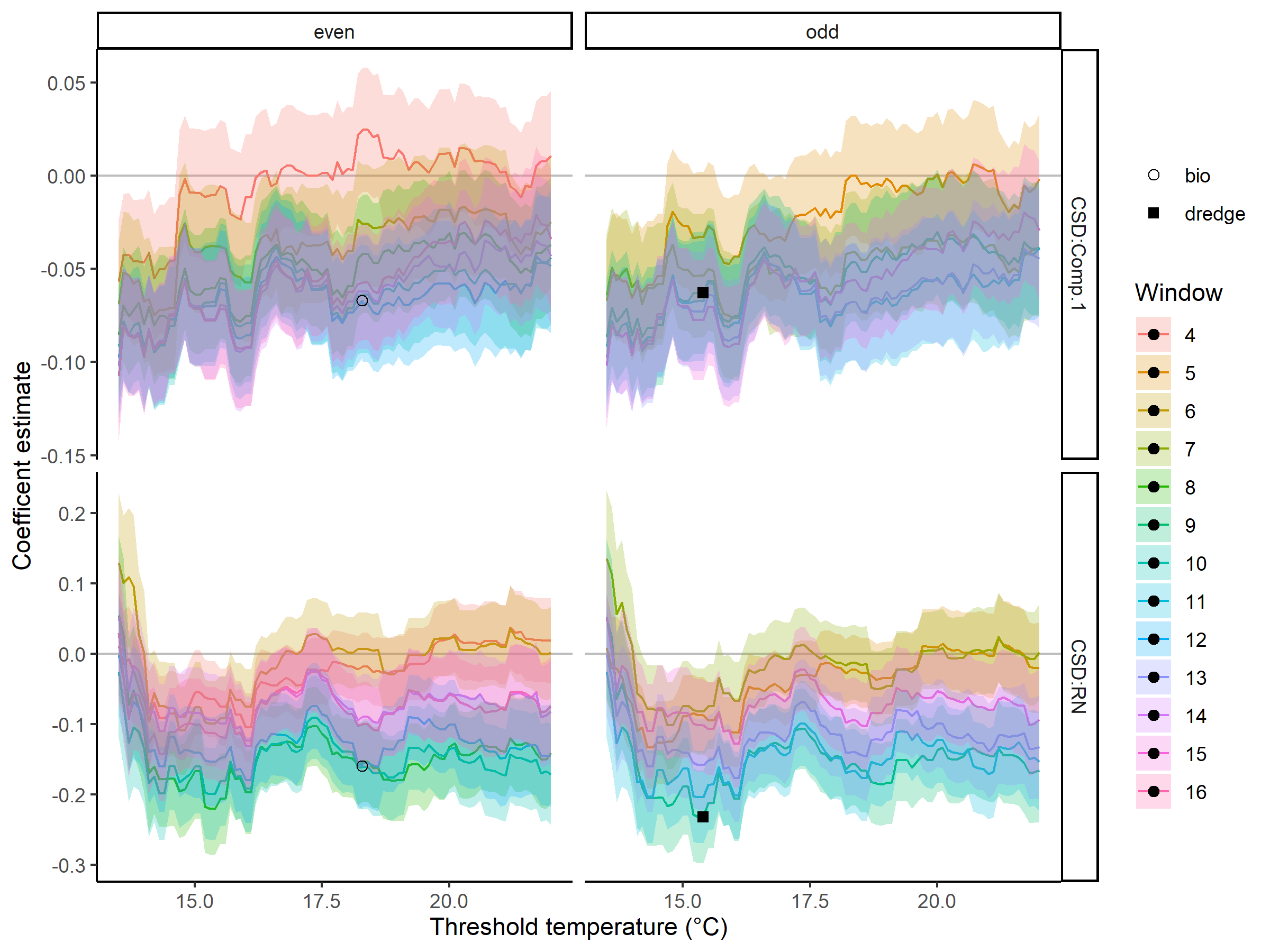

**Fig. S3:** Post hoc dredging analysis assessing the optimal threshold temperature determining fledging success. Presented are the coefficient estimates and 95% CI for the main interaction terms of interest with threshold temperatures increasing by 0.1°C. Presented colors are the post-hatching widow increasing between 4 and 16 days post hatching. The two points represent: the combination used in the text body (bio) and the combination of the model with the lowest AICc (dredge). The two panels are the figure separated by if the post hatching window was either an odd or evenly numbered window.

**Differing definition of cold snaps and precipitation**

To further evaluate the suitability of defining cold snaps as days in which the maximum temperature fell below the 18.3°C temperature threshold, we evaluated a series of other weather-related definitions and how they influenced the predictability of Tree Swallow fledging success. We first defined a cold snap as a day in which the maximum (Max), minimum (Min) and mean (Mean) temperature fell below the 18.3°C threshold. Cold snaps for each brood were then defined as the number of days below these temperatures during the 12-day post hatching window. We included another definition of temperature (Mean daily max), defined as the mean daily maximum temperature observed during the 12-day post hatching window. We further explored two precipitation definitions. First the mean daily precipitation and second the number of days in which precipitation of greater than 2 ml (median observed value) was recorded during the 12 day post hatching window. We then integrated these definitions into identical models (Land * Temp in Appendix S2; Table S1) and evaluated their performance via AICc values (Table S1). In all cases, our original definition far outperformed that of any other definition.

**Table S1**: The results of a model comparison in which identical models (Land * Temp in Appendix S2; Table S1) received differing definitions of the number of cold snap days and precipitation. Presented are all the combinations of possible weather definitions and their model’s AICc values represented as the delta from the model with the lowest AICc. The model with no information for either temperature or precipitation is a model without these terms.

| Temperature definition | Rain group | ΔAICc | *w* |
| --- | --- | --- | --- |
| Max | Daily value | 0.000 | 1 |
| Max | Greater than >2 ml | 17.343 | 0 |
| Mean | Daily value | 147.832 | 0 |
| Mean | Greater than >2 ml | 165.594 | 0 |
| Min | Daily value | 87.542 | 0 |
| Min | Greater than >2 ml | 106.091 | 0 |
| Mean daily max | Daily value | 67.646 | 0 |
| Mean daily max | Greater than >2 ml | 89.369 | 0 |
|  |  | 180.750 | 0 |

**Literature cited.**

Burnham, K. P., and D. R. Anderson. 2002. Model Selection and Multimodel Inference. second. Springer.

Cade, B. S. 2015. Model averaging and muddled multimodel inferences. Ecology 96:2370–2382.

Glądalski, M., M. Bańbura, A. Kaliński, M. Markowski, J. Skwarska, J. Wawrzyniak, P. Zieliński, and J. Bańbura. 2020. Extreme temperature drop alters hatching delay, reproductive success, and physiological condition in great tits. International Journal of Biometeorology 64:623–629.

Harrison, X. A., L. Donaldson, M. E. Correa-Cano, J. Evans, D. N. Fisher, C. Goodwin, B. Robinson, D. J. Hodgson, and R. Inger. 2017. Best practice in mixed effects modelling and multi-model inference in ecology. PeerJ Preprints 5:e3113v1-48.

Roitberg, B. D., and M. Mangel. 2016. Cold snaps, heatwaves, and arthropod growth. Ecological Entomology 41:653–659.

Taylor, L. R. 1963. Analysis of the Effect of Temperature on Insects in Flight. The Journal of Animal Ecology 32:99–117.

Winkler, D. W., M. K. Luo, and E. Rakhimberdiev. 2013. Temperature effects on food supply and chick mortality in tree swallows (Tachycineta bicolor). Oecologia 173:129–138.

de Zwaan, D. R., A. F. Camfield, E. C. MacDonald, and K. Martin. 2019. Variation in offspring development is driven more by weather and maternal condition than predation risk. Functional Ecology 33:447–456.

de Zwaan, D. R., A. Drake, J. L. Greenwood, and K. Martin. 2020. Timing and Intensity of Weather Events Shape Nestling Development Strategies in Three Alpine Breeding Songbirds. Frontiers in Ecology and Evolution 8.
