## Appendix S4 for "Interacting effects of cold snaps, rain, and agriculture on the fledging success of a declining aerial insectivore"

### Appendix S4: Supplemental analysis of cold snap appearances and variation in hatching date

### Likelihood to encounter a cold snap

Because of the substantial longitudinal and elevational change expressed by the study system, we investigated if the likelihood to encounter a cold snap varied throughout the various breeding seasons and along the agricultural gradient. We therefore modeled the daily likelihood of experiencing a cold snap throughout the study of fledging success. We used the daily temperatures collected on each farm between the years of 2006 and 2016 between 1 June to 15 July (N = 19,800 day/farm/year). We then assessed if the maximum temperature of each day on each farm was either above the critical threshold (no cold snap = 0) or below the critical threshold (snap = 1). We constructed two models both evaluating the daily likelihood of experiencing a cold snap using GLMMs with a binomial distribution, and because we observed the probability of a cold snap was low, used a Complementary Log-Log (c-loglog) link function. The first model included the year, Julian date, their interaction as fixed effects and the farm ID as a random effect. We observed that there was a slight increase in the daily likelihood of a snap to occur throughout the data set (Table S1 and Fig. S1). The second model included the Julian date and both site scores as fixed effects and the year as a random effect. We found that indeed the daily likelihood of experiencing a cold snap varied with landscape context, and that broods within more forage landscapes were almost twice as likely to experience at least one cold snap than any other landscape context (Table S2 and Fig. S2).

**Table S1**: Results of modeling the daily likelihood in the occurrence of a cold snap from 19,800 daily observations across 40 separate farms and 11 years along a gradient of agricultural intensification. Models treated the appearance of cold snaps as a binomial response with 1 being the appearance of a cold snap and 0 being none. Models included the interaction between the Julian date of temperature recording and the year as fixed effect and the farm as a random effect.

| Term | Beta | 95% CI^1^ | p-value |
| --- | --- | --- | --- |
| Julian date | -1.2 | -1.3, -1.1 | <0.001 |
| Year | 0.42 | 0.33, 0.50 | <0.001 |
| Julian date * Year | 0.39 | 0.32, 0.46 | <0.001 |
| ^1^CI = Confidence Interval |  |  |  |

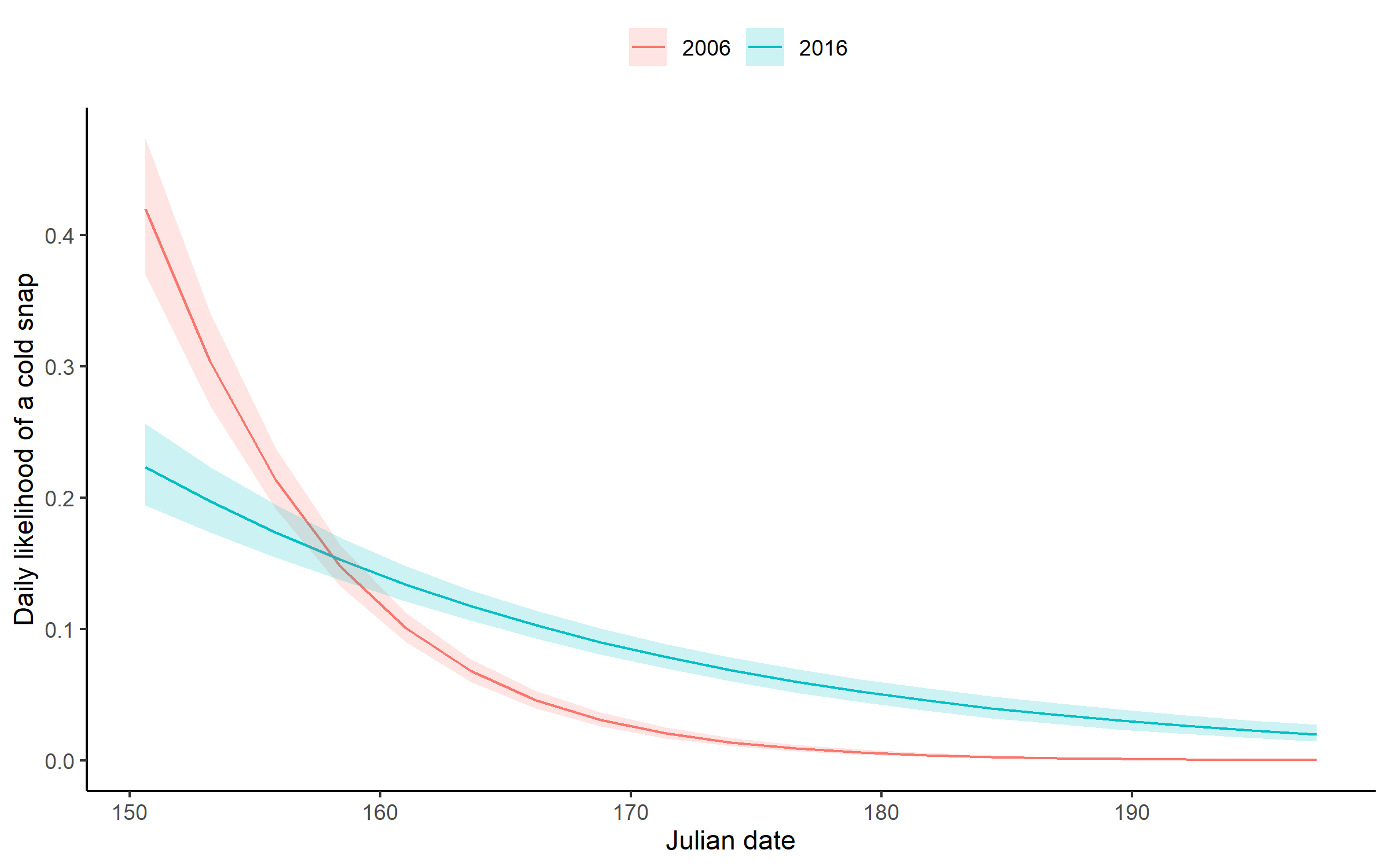

**Fig. S1 :** Predictions of daily likelihood of a cold snap occurring as a function of the interaction between Julian date and the breeding season. Models treated the appearance of cold snaps as a binomial response with 1 being the appearance of a cold snap and 0 being none. Models included the interaction between the Julian date of temperature recording and the year as fixed effect and the farm as a random effect.

**Table S2:** Standardized coefficient estimates and their 95% confidence intervals evaluating the likelihood of experiencing a cold snap during our study on the fledging success of Tree Swallows across a gradient of agricultural intensification within southern Québec, Canada, between 2006 and 2016 (N = 19,800). Fixed effects include the Julian date of temperature recording, site scores, and their interaction. The GLMM with binomial response and c-loglog link function included year and farm IDs as random effects.

| Term | Beta | 95% CI^1^ | p-value |
| --- | --- | --- | --- |
| Julian date | -1.3 | -1.3, -1.2 | <0.001 |
| Comp.1 | -0.07 | -0.10, -0.03 | <0.001 |
| Comp.2 | -0.07 | -0.14, -0.01 | 0.034 |
| Comp.1 * Comp.2 | 0.06 | 0.02, 0.09 | 0.003 |
| ^1^CI = Confidence Interval | |  |  |

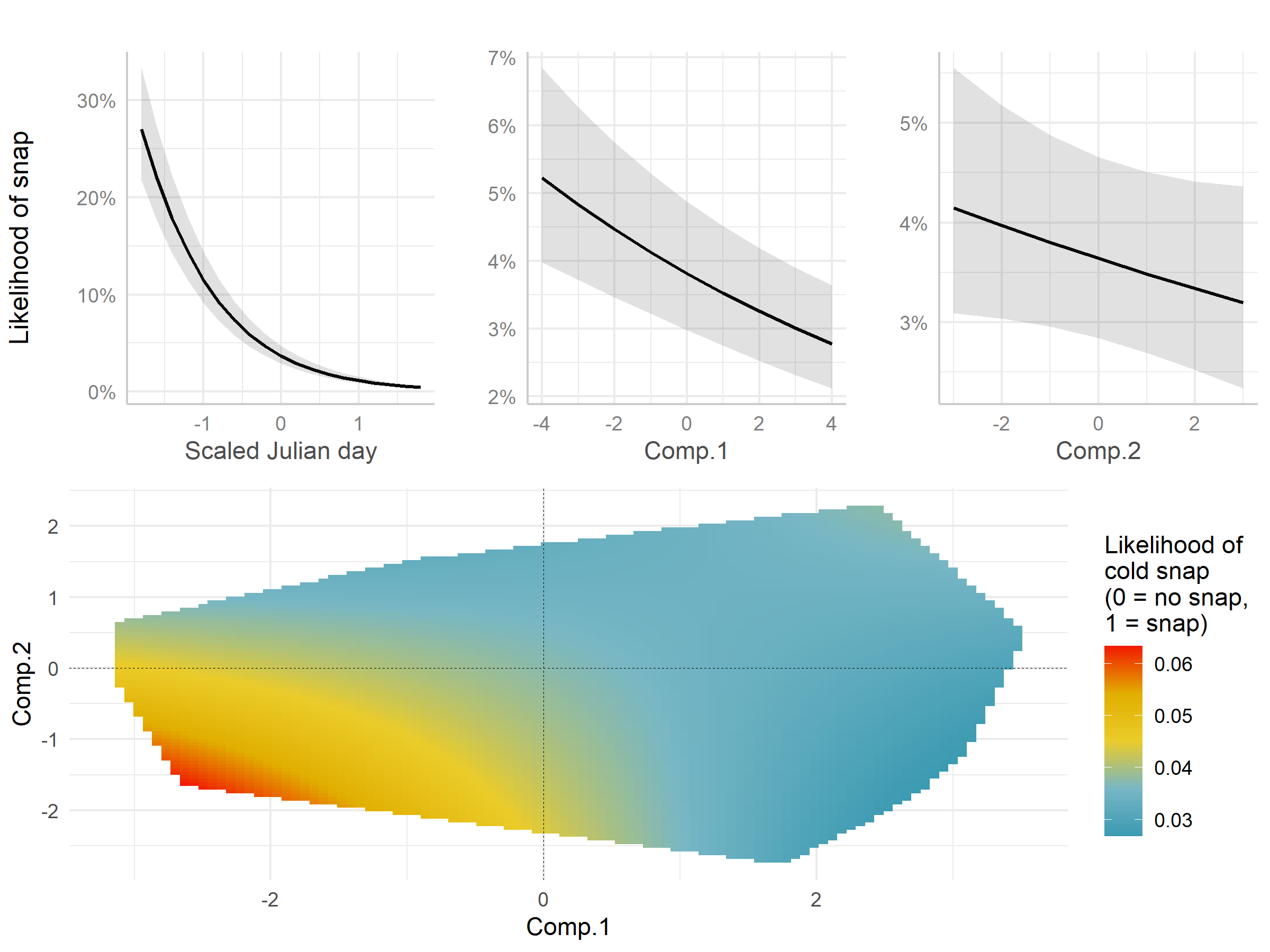

**Fig. S2:** Predictions of the daily likelihood of experiencing a cold snap against the scaled Julian date as well as the first two components deriving landscape context.

### Advancing hatching dates

We desired to evaluate the hypothesis that, as a response to climate change, migratory birds may advance the onset of their breeding attempts using our data set. We therefore modeled the hatching date of broods as a function of the breeding season using linear mixed effects models. We included year as a continuous fixed effect and the farm and breeding female ID as random effects. We observed that indeed breeding events have advanced by nearly 1.1 days in the dataset used for this analysis (Table S3, Fig. S3).

**Table S3:** Standardized coefficient estimates and their 95% confidence intervals evaluating the if hatching dates of Tree Swallows has advanced during a study on their fledging success across a gradient of agricultural intensification within southern Québec, Canada, between 2006 and 2016 (N = 1,897). The LMM included hatching date as a fixed effect and year and farm IDs as random effects.

| Term | Beta | 95% CI^1^ | p-value |
| --- | --- | --- | --- |
| Year | -0.11 | -0.19, -0.03 | 0.007 |
| ^1^CI = Confidence Interval | | | |

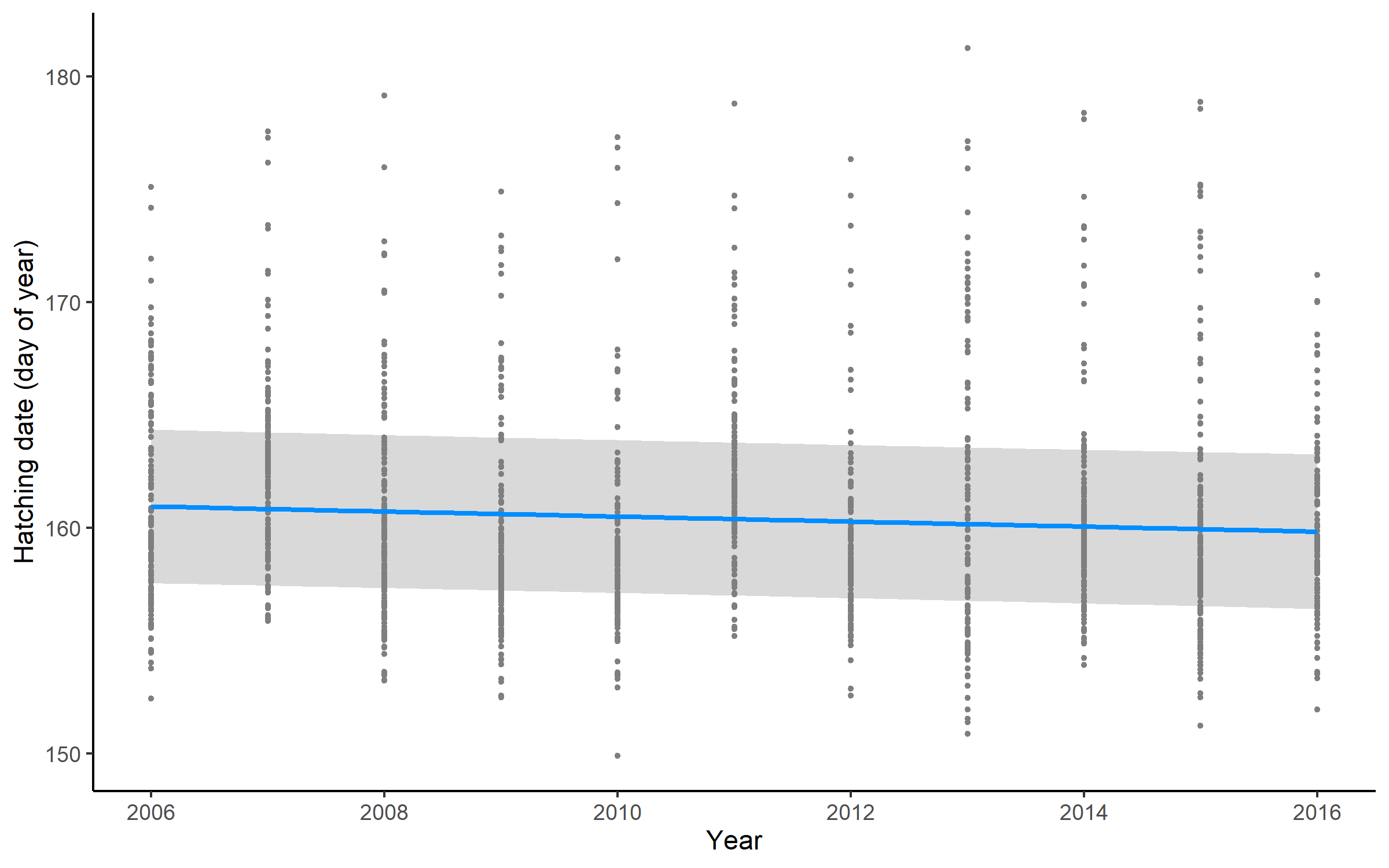

**Fig. S3:** Predictions and 95% confidence intervals of hatching date against breeding season. Points in background are partial residuals. The LMM included hatching date as a fixed effect and year and farm IDs as random effects.
